# First study of an aye-aye (*Daubentonia madagascariensis*) mother’s anxiety behaviors peripartum

**DOI:** 10.1101/2025.11.08.686247

**Authors:** Rebecca V. Olson, David Watts, Lisa M. Paciulli

**Affiliations:** Department of Cell Biology, University of Virginia, Charlottesville, VA, USA; Research PackTrack Program, Department of Biological Sciences, North Carolina State University, Raleigh, NC, USA; Federal Law Enforcement Training Center (FLETC), GA, USA; Department of Biological Sciences, North Carolina State University, Raleigh, NC, USA

**Author notes:** **Corresponding Author** Rebecca V. Olson University of Virginia, Department of Virginia.

## Abstract

Anxiety is a negative psychological state induced by stress triggers that can be examined in animals which exhibit repetitive, self-directed behaviors. Primate mothers face anxiety due to birthing, infant health, and social relations. In this study, a mother aye-aye (*Daubentonia madagascariensis*) was observed to assess anxiety behaviors. It was hypothesized that the behaviors would change over time. With Duke IACUC approval, Pelco IMM12027-1S cameras were placed in the Duke Lemur Center (DLC) aye-aye mother’s enclosure to record her activity. Over three days peripartum, full 24-hour footage and continuous focal animal sampling were used to note the time, duration, and type of repetitive scratching, grooming, etc. Data were then analyzed using Chi-square, Fishers Exact, and Kruskal Wallis tests. The results showed that the day before birth, the mother repeatedly constructed and deconstructed nests eight times more than engaging in any other behavior (n=212x), while on the day of birth, the mother scratched (n=87x) and groomed (n=60x) herself a lot. The day after birth, the mother was hyper-vigilant (n=32x). Thus, the hypothesis was supported as the mother exhibited signs of anxiety by engaging in different anxiety-related behaviors over the three-day period. While limitations such as a small sample size (n=1) and possible abnormal parturition behavior are evident, this study is the first to examine maternal anxiety in aye-ayes. The results can help husbandry staff create more comfortable environments for the rare and difficult-to-see captive aye-ayes.

## Introduction

Anxiety affects 359 million people worldwide (Global Burden of Disease, 2021) and is defined as a negative psychological state induced by stress triggers (Coleman & Pierre, 2014). Stress is an individual’s response to a pressure that threatens homeostasis and requires the individual to adapt (Del Giudice et al., 2018, Moberg & Mench, 2000: 3, Antelman & Caggiula, 1980). In this paper, the anxiety behavior of a mother aye-aye lemur (*Daubentonia madagascariensis*) peripartum is examined.

In humans, stress can be caused by external and internal factors such as work, school, family pressures, financial issues, relationship problems, major life changes, and feelings overwhelm (Elias, 1989; Bhargava & Trivedi, 2018; Short, 2002; Ryu & Fan, 2023; Harb et al., 2024; Downey & Crummy, 2022 ; Kendler et al., 1999; Finlay-Jones & Brown, 1981; Wang & Saudino, 2011; Chu et al., 2024). In nonhuman animals, stress triggers include conflicting drives, fear, social and environmental stressors, aggression, and unachievable goals (Maestripieri et al., 1992). Physiological responses to stress include activation of the sympathetic nervous system and include agitation, sweating, and an increased heart rate (Coleman & Pierre, 2014). People who are anxious act irritable, nervous, and restless (Coleman & Pierre, 2014).

Related to stress is anxiety, the behavioral, psychological, and physiological state of heightened arousal (Steimer, 2002). Anxiety likely evolved as a survival mechanism to warn an individual of danger and to trigger adaptive responses to bring the body back to homeostasis (Marks & Nesse, 1994; Antelman & Caggiula, 1980). Increased survival behaviors like threat assessment and cooperative behaviors can promote resiliency in stressful or threatening environments (Gallup, 2022; Palagi et al. 2009; Maestripieri et al., 1992).

There has been a rise in maternal mental health and anxiety-related disorders in human primate mothers (Daw et al. 2025; McKee et al. 2020). The anxiety and stress of being a mother and having to be responsible for offspring survival can cause and/or trigger anxiety (Jeličić et al. 2022). Thus, mothers commonly face anxiety peripartum especially immediately before and/or after the birth of a child (Radoš et al. 2018). The anxiety and stress felt by mothers is likely related to fetal motility, heart rate, and behavior, as well as concerns regarding delivery, infant health, and social relations (Radoš et al., 2018). A mother’s response to these triggers can be heightened postpartum due to hormonal shifts and physical discomfort, changes in routine, and the stress of caring for a new infant, all of which can increase anxiety (Ballard et al., 1993). In a study on postpartum depression and anxiety in people, Radoš et al. (2018) found that mothers faced the highest levels of anxiety in their last trimester of pregnancy and in the two days following birth.

Mother-infant separation is also a main trigger of anxiety. Maternal separation anxiety is the unpleasant emotional state that mothers, in particular, face when being separated from their infant (Hock et al., 1989). Both intrinsic traits, such as genetically determined biases, and experiential factors, such as previous relationship histories and social structure, can heighten separation anxiety in mothers (Hock et al., 1989). The bonding between mother and infant induces a strong attachment, which additionally influences maternal anxiety (Karayağız et al., 2020). Parental bonding is formed early and is strengthened as the infant matures and interacts with the maternal individual (Karayağız et al., 2020).

In mothers, anxiety can also increase responsiveness to threats as well as maternal attentiveness to, and affiliative behavior toward, their infants (Maestripieri, 2005, Nguyen, 2008). Maternal anxiety may also help the infant respond better to stressful postnatal environments by predisposing them to challenges (Dantzer et al., 2013; Love et al., 2013; Bauer et al., 2019; Gluckman & Hanson 2004; Lee & Zucker 1988; Barker 2002). That said, evolved behaviors are generally context-dependent and thus, while maternal anxiety can be beneficial in some sense, it may also have some negative consequences such as suppressing maternal motivation and enhancing rejection of an infant (Sanchez, 2010; Bardi et al., 2004; Saltzman & Abbott, 2009; Maestripieri et al., 2009).

It only makes sense that nonhuman animals also suffer from anxiety. Animal anxiety often takes the form of self-directed actions or behaviors (SDBs) and/ or stereotypic behaviors / stereotypies (STBs) (Maestripieri et al., 1992; Troisi, 2002). SDBs are actions an individual performs on their own body such as auto (self)-touching, auto (self)-scratching, hair picking, and auto (self)-grooming (Maestripieri et al., 1992). These behaviors are similar as self-touching is defined as brief contact with fingers and other parts of the body, without scraping or picking (Castles et al., 1999; Laméris et al., 2022), while autoscratching is when the individual scrapes their claws or nails against its skin (Easley et al., 1987, Aureli & van Schaik, 1991). Hair picking is the act of repeatedly pulling or removing hair from the body (Woods & Houghton, 2014), and auto-grooming is using the hand, feet, or mouth to move, sweep, and/or rub against the fur, facial orifices, and/or genital region (Troisi and Schino, 1987, Lopez-Vergara et al., 1989).

SDBs typically occur in response to stressful events such as social conflict and reflect heightened physiological arousal (Troisi, 2002). These behaviors may be adaptive, functioning to help the individual prepare themselves for a threat or negative interaction (Maestripieri et al., 1992).

However, when SDBs become repetitive and chronic, they can develop into stereotypic behaviors/ stereotypies. While STBs may include some SDBs, namely those directed at the self, they are more broadly defined as rhythmic, repetitive, and invariant behaviors that lack an obvious goal or function (Mason, 1991). Additional examples include yawning, panting, shaking, and/or pacing (Maestripieri et al., 1992; Schino et al., 1996; Aureli et al., 2002, Lutz et al., 2003). Yawning is involuntarily opening the mouth (Gupta & Mittal, 2013), and panting is when the individual displays open mouth respiration characterized by moderate to rapid airflow (Schmidt-Nelsen et al., 1970; Goldberg et al., 1981). Shaking is when an animal trembles in place (Maestripieri et al., 1992; Aureli & van Schaik, 1991), while pacing is continuously moving, such as jumping or traveling, back and forth between an area with a fixed pattern (Cless et al., 2015; Burgener et al., 2008; Lutz et al. 2003; Berkson, 1968).

STBs can arise when animals experience boredom or environmental deprivation in captivity (Barnett & Hemsworth, 1990; Fernandez, 2021) and/or are provoked by stressful or anxiety-inducing stimuli (Maestripieri et al., 1992). Unlike SDBs, stereotypies are more often associated with negative, maladaptive states and may reflect physiological and behavioral issues (Wielebnowski, 2003). Even the location of these behaviors may reveal their underlying cause. For example, anxiety-related, stereotypic behaviors are common in captivity yet rarely observed in natural habitats (Carlstead, 1998). Further differentiating SDBs and STBs, a study on red-capped mangabeys (*Cercocebus torquatus*) revealed that individuals housed in both a captive and a free-range settings displayed SDBs, but the captive individuals also displayed high prevalence of maladaptive STBs like pacing (Reamer et al., 2010). This potentially reflects abnormal behavior and interactions with abnormal environments (Carlstead, 1998). Similarly, sleep-deprived captive chimpanzees (*Pan troglodytes*) have been observed performing stereotypies in their nests (Berkson, 1967).

STBs are also thought to serve as coping mechanisms that help the individual maintain psychological stability (Fox 1984, p. 184). They may have a soothing or self-calming effect, allowing the release of endorphins (Tatemoto et al., 2022; Salali et al., 2021). It is likely that the release of the "feel-good" chemicals helps relieve the body of pain and promote a sense of well-being (Simmons & Self, 2009; Matthew & Paulose, 2011). Bank voles (*Clethrionomys glareolus*) injected with naloxone, an opioid antagonist, and who were put in stressful situations such as barren laboratory cages, showed a reduction in STBs (Kennes et al., 1988), further showing that stereotypies function as self-coping mechanism.

Both STBs and SDBs are widely used as noninvasive, reliable behavioral indicators of anxiety (Maestripieri et al., 1992; Schino et al. 1996; Aureli et al., 2002; Troisi, 2002). Moving forward in this paper SDBs and other behaviors typically categorized as STBs like (yawning, panting, shaking, and/or pacing) will be collectively referred to as “anxiety-related behaviors”. It should be of note that anxiety-related behaviors can only be considered stereotypic when they meet the criteria of being chronic, repetitive, invariant, and functionally associated with coping and/ or as a maladaptive response.

### Nonhuman Primate Anxiety

Nonhuman primates (lemurs, monkeys, and apes) exhibit many anxiety behaviors, and may engage in STBs when anxious (Troisi, 2002; Baker & Aureli, 1997; Marriner & Drickamer, 1994; Marais et al., 2006). For example, after intragroup aggression, captive long-tailed macaques (*M. fascicularis*) exhibit several anxiety-related behaviors. These include the victim shaking, as well as auto-scratching and auto-grooming (Aureli et al., 1989; Aureli & van Schaik, 1991). Electrical and pharmacological activation of the locus coeruleus, the major brain noradrenergic nucleus that controls autonomic functions such as the stress response, also elicited scratching and other behaviors in stump-tailed macaques (*M. arctoides*) similar to those observed in monkeys exposed to natural threats in the wild (Redmond & Huang, 1979; cf. Troisi et al., 1991). Ninan et al. (1982) administered fl-CCE to increase feelings of anxiety to rhesus macaques (*Macaca mulatta*). The anxiogenic substance produced a behavioral response similar to fear or anxiety in the macaques who also scratched more. Also, auto-scratching rates in ring-tailed lemurs (*Lemur catta*) increase during estrus. This is most likely due to hormonal changes, intersexual competition, and mating unpredictability (Sclafani et al., 2012). Auto-scratching in wild brown lemurs (*Eulemur fulvus*) was found to increase with intergroup aggression and predation attempts (Palagi & Norscia, 2010). All of this provides evidence for the link between scratching behaviors and stress.

Repeated hair-picking has been observed in baboons (*Papio anubis*), rhesus macaques (*M. mulatta*), and chimpanzees (*P, troglodytes*). Females are more prone to picking their hair than males (Reinhardt, 2005). In addition, increased rates of auto-grooming and other potential stereotypies like yawning, shaking, and scratching were seen in free-ranging wild vervet monkeys (*Chlorocebus aethiops*) (McDougall, 2011). When vervets were in proximity to more dominant and/or nonassociated conspecifics, they were more likely to exhibit anxiety-related behaviors (McDougall, 2011).

Anxious captive rhesus macaques (*M. mulatta*) have been observed pacing especially during or after acute stressors. For example, macaques paced when they were moved out of their home-cage, when they were being transported, and when they were exposed to a conditioned cue of imminent separation from their partner (Mitchell & Gomber, 1976; Willott & McDaniel, 1974). Poirier et al. (2019) found that macaque pacing could also be a result of chronic stressors, such as the agonistic behavior of conspecifics in a nearby enclosure (Poirier et al. 2019).

Stump tailed macaques (*Macaca arctoides*) yawn when watching videos of unfamiliar monkeys yawning, potentially alluding to the evolution of enhanced vigilance against threats and as a way to foster a cooperative environment (Paukner & Anderson, 2006; Gallup, 2022; Palagi et al., 2009). When the macaques yawned, they also scratched themselves, indicating an increased level of anxiety (Paukner & Anderson, 2006). Also, a positive correlation between yawning and auto-grooming was found in gelada baboons (*Theropithecus gelada*) (Palagi et al., 2009). This suggests that when engaged in an anxiety-related behavior such as auto-grooming, gelada baboons were more likely to be triggered into an additional stress response such as yawning (Palagi et al., 2009).

Thus, when nonhuman primates feel anxious and stressed, they engage in anxiety-related behaviors such as auto-grooming, auto-scratching, hair picking, pacing, and yawning (Maestripieri et al., 1992; Schino et al., 1996; Aureli et al., 2002).

### Primate maternal anxiety

Mothering and keeping another individual alive is a stressful business (Human: Fleming et al., 1997; rodents: Deschamps et al., 2003; nonhuman primates: Brent et al., 2002 Bardi et al., 2004; Hoffman et al., 2010; Troisi et al., 1991). Mothers have extra stressors to contend with before, during, and after birth and interactions with infants are the main source of anxiety for mothers (Troisi et al., 1991).

Maternal anxiety in nonhuman primates can also be measured through anxiety-related behavior. Captive black-and-white ruffed lemur (*Varecia variegata*) mothers spent 75% of their time with their infant in the first few weeks of life, which helps enhance the parental bond (Klopfer & Dugard, 1976). However, when initially separated from one another, both the mother and the infant showed high levels of anxiety. During such separation events, ruffed lemur mother-infant pairs auto-groomed more, indicating an increase in anxiety (Klopfer & Dugard, 1976).

Captive Japanese macaque (*Macaca fuscata*) maternal anxiety was examined in regard to scratching prior to and post-birth (Troisi et al., 1991). Troisi et al. (1991) found that macaque mothers scratched themselves more postpartum regardless of their rank in the dominance hierarchy. The results showed that behavioral anxiety in mothers after birth is dependent upon maternal stressors and not just social pressures. The rate of scratching in macaques was also evaluated by Maestripieri (1993), who found that scratching increased when an infant was separated from its mother. Maternal scratching also increased when an infant was approached by an individual who had previously harassed them and when in spatial proximity to an adult male or high-ranking female (Maestripieri, 1993a). These results show that auto-scratching is an indicator of anxiety in mother macaques and is especially high during the birthing season and potentially dangerous situations reflecting the mothers states of uncertainty (Maestripieri, 1993a).

Examination of genitalia is the inspection of the genital region without hair picking, auto-grooming, or auto-scratching of the region (Ocepek & Andersen, 2018; Turner et al., 2010). This behavior has been observed in free ranging Japanese macaques and captive long-tailed macaques (*Macaca fascicularis*) where genital examination followed by the sucking or licking of their hands increased in frequency around the time of birth (Kemps & Timmermans, 1982; Turner et al., 2010). Although these behaviors have not been directly linked to anxiety, they are classified as a self directed behavior, which may indicate anxiety in primates (Maestripieri et al., 1992, Schino et al., 1996, Aureli et al., 2002, Troisi, 2002).

Vigilance is hyper-alertness accompanied with widened and opened eyes, the head angled up, and remaining still in fixation (Kutsukake, 2006). In a study on captive rhesus macaque vigilance (*M. mulatta*), the rate of a mother’s vigilance over her infant continued to increase after the birth (Maestripieri, 1993b). This was most likely due to additional maternal stressors such as a newborn’s inability to move around much and not being safe independently. Rhesus macaque vigilance behaviors when the infant was separated from the mother were more frequent in the first few weeks of an infant’s life and remained high when the infant was not near the mother, regardless of the infant’s age (Maestripieri, 1993b). Maestripieri interpreted increased visual monitoring of an infant as reflecting the mothers’ fear component of maternal anxiety.

Repeated nest construction is the repetitive manipulation of browse, branches, leaves, blankets, paper, etc. in the nest in order to construct a comfortable environment (Goodall, 1968, Kano, 1979, Fruth & Hohmann, 1996, Feistner & Ashbourne, 1994). Preparing for birth, a wild chimpanzee (*P. t. schweinfurthii*) mother built three nests in the span of two hours before parturition. She also built two new nests immediately after giving birth (Goodall & Athumani, 1980). The mother chimp was also observed to be very restless.

Excessive fussing over an infant and constantly touching or moving them could indicate maternal stress. We called this behavior “nudging” and defined it as when a mother repositioned the infant using her nose, hands, or trunk, and during this time, the mother does not provide direct care (such as grooming) to the infant (Silk et al., 2003; Cheney & Seyfarth, 1999; Dunayer and Berman, 2018). Schino et al. (2003) found that lower ranking, younger captive Japanese macaque (*M. fuscata*) mothers physically handled their infants in a more protective manner and that these mothers were often more anxious (more in macaques: Maestripieri, 1993a; Maestripieri, 2001). Anxious rhesus macaque (*M. mulatta*) mothers may maintain high contact with their offspring and restrain them from moving away, exploring, and gaining independence especially in stressful or threatening events (McCormack et al., 2015). Also, Hooley & Simpson (1981) regarded maternal anxiety as the factor causing primiparous rhesus mothers to hold and restrain their infants physically more than did the multiparous mothers.

Thus, nonhuman primate mothers are stressed before, during, and for at least several weeks after giving birth - if not forever. Primate mothers reveal their levels of anxiety with general restlessness and increased self-grooming (Troisi et al. 1991; Klopfer & Dugard, 1976, Feistner & Ashbourne, 1994), self-scratching (Troisi et al., 1991, Maestripieri, 1993a), and repeated examination of their genital region (Kemps & Timmermans, 1982; Turner et al., 2010). Additionally, increased vigilance (Maestripieri, 1993b), repeated nest-building (Goodall & Athumani, 1980), as well as and nudging of their offspring (Schino et al., 2003) may indicate maternal anxiety.

While there are many studies on anxiety behavior in people (see Mishra & Varma, 2023 for a recent review) and in human mothers (Review: Araji et al., 2020), little has been published on anxiety behavior in aye-ayes (although see Bass 2016 for an exception) and even less has been written about aye-aye maternal anxiety. Therefore, in this study, anxiety behavior in a mother aye-aye (*Daubentonia madagscariensis*) was examined. It was hypothesized that the mother aye-aye would exhibit behavioral anxiety peripartum and that the anxiety would differ before and after birth.

## Methods

### Study Subject and Area

The subject of this study was a Duke Lemur Center aye-aye (*D. madagscariensis*) mother. Aye-ayes are the world’s largest nocturnal primate, naturally occurring in Madagascar (Kay & Kirk, 2000). Aye-ayes are in the family Daubentoniidae, they are Endangered (Louis et al., 2020; IUCN 2025), and have several unique characteristics such as large incisors, ears, and eyes, and a long middle finger for percussive foraging (Soligo, 2005; Erickson, 1991). Aye-ayes also have a large brain size to body size ratio (1:70: Kaufman et al., 2005) as well as a pseudothumb (Hartstone-Rose et al., 2020).

Aye-ayes build cocoon-shaped nests 10-15m above the ground with small branches and vines (Petter & Peyrieras, 1970, Quinn & Wilson, 2004). The nests are closed except for a 15cm opening (Petter & Peyrieras, 1970). While adult aye-ayes spend most of their time alone, aye-aye mothers spend 82% of their time with their newborns (Feistner & Ashbourne, 1994). Infants sleep in the same nest as their mother (Petter & Peyrieras, 1970), and may cling to their mothers for one to two years after birth (Feistner & Ashbourne, 1994; Rakotondrazandry et al., 2021). Feistner and Ashbourne (1994) observed that a captive, almost one-year-old aye-aye occasionally slept in a separate nest from the mother. It takes infant aye-ayes up to two years to become independent because they need time to develop the fine motor skills necessary for foraging and survival on their own (Feistner & Ashbourne, 1994; Rakotondrazandry et al., 2021).

Some anxiety-related behaviors have previously been observed in aye-ayes and include pacing, hyper-vigilance, and over-grooming (Bass, 2016). Aye-ayes paced more and showed increased vigilance when they were observed by someone in person compared to when no human was present and the aye-ayes were observed in video-recordings. The pacing and vigilance was associated with increased levels of cortisol (Bass, 2016). Hyper-alert aye-ayes also perk their ears, increasing auditory vigilance (Owen, 1866; Erickson, 1991, 1995). In addition, when a captive infant was not near their mother two weeks postpartum, the mother aye-aye self (auto)-groomed more. This indicated higher levels of anxiety (Feistner & Ashbourne, 1994). There are approximately 25 aye-ayes living in the United States (Duke Lemur Center, 2016). Most of the aye-ayes (∼10) are housed at the Duke Lemur Center (DLC) in Durham, North Carolina (35.9941° N, 78.9604° W) (Duke Lemur Center, 2016).

The DLC aye-ayes live under a reverse light cycle so they are active during the day when people are around (E. Ehmke, pers. comm.). The aye-aye mother (*Medusa*, DOB: 09/14/2003) lived in an enclosure with cinder block walls arranged in a hexagonal form. The enclosure was 17.5 feet (5.3 meters) long by 15.2 feet (4.6 meters) wide and 16.2 feet (4.9 meters) tall with a volume of approximately 3,230 cubic feet (pers. obs.).

### Data Collection

With Duke Institutional Animal Care and Use Committee approval (IACUC A202-16-09), DW placed three Pelco IMM12027-1S motion-sensor cameras around the mother’s (*Medusa*) enclosure before she was expected to give birth. One camera was placed in the center of the enclosure and one camera was put in each of the two nest boxes (15.75"x 16"x 18" stainless steel) so that the mother could be seen at all times. **Figure 1** shows the enclosure and placement of the cameras. The cameras were set to video-record the female’s behavior 24 hours a day, and the resulting videos were uploaded to Google Drive. Medusa gave birth to a female (*Agatha*) at approximately 1:50 a.m. on June 7, 2017. Thus, Medusa’s behavior was captured on video the day before (June 6), the day of (June 7), and the day after she gave birth (June 8). In this paper, Medusa’s anxiety behavior peripartum - around the time of birth - was examined.

**Figure 1:**
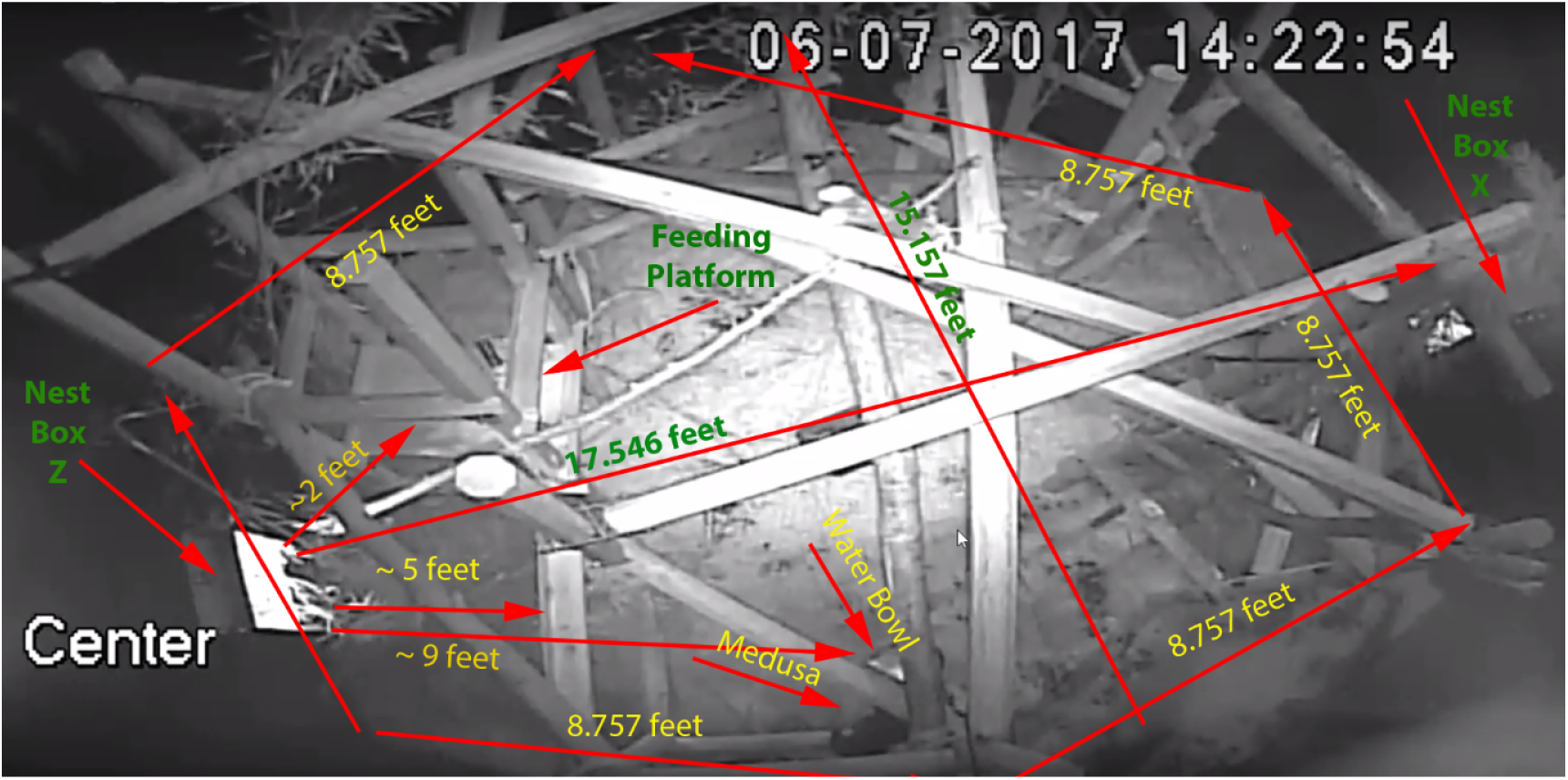
Photo of mother Medusa aye-aye’s enclosure showing dimensions. Nest Box Z is on the left side of the enclosure, while Nest Box X is on the right side. Medusa can be seen in the bottom middle of the photo twelve hours after giving birth.

Paciulli Laboratory undergraduate research assistants (RA’s) were trained how to identify aye-aye behaviors in sample training video files. RA’s had to pass the video file training with 100% accuracy before having access to the peripartum video footage. RA’s used all occurrences focal animal sampling methodology (Altman, 1974; Martin & Bateson, 2007) to identify anxiety-related behaviors and potential stereotypies. Anxiety behaviors included auto(self) grooming, auto(self) scratching, examining genitalia, hair picking, hyper-vigilance (defined as observing on coder observation forms), nest construction, nudging infant, pacing, panting, shaking, and yawning. See **Table 1** for behaviors and operational definitions.

**Table 1:** Operational definitions of anxiety-related behaviors examined in *Medusa*, the mother aye-aye.

| Behavior | Definition |
| --- | --- |
| Auto (self) Grooming | The individual uses their hand, feet, or mouth to rub, lick, and/or sweep against their fur, genital region, or facial orifices. This behavior is associated with cleaning and general maintenance of the body |
| Auto (self) Scratching | Scraping of an individual's claws or nails against its skin. |
| Examining Genitalia | The individual inspects the genital region without participating in hair picking, auto-grooming, or auto-scratching of that region. |
| Hair Picking | Repeated removal or pulling of hair from the body which may lead up to significant hair loss. |
| Hyper-vigilance (Observing) | The individual remains still in fixation, widened and opened eyes, and gaze focused on their surroundings. Ears are also perked. |
| Nest Construction | The manipulation of browse, blankets, paper, etc. in the nest box in order to build a cocoon like nest. |
| Nudging infant | The repositioning of the infant by using the nose, hands, or trunk.<br>The individual does not provide direct care or discernable benefit to the infant or herself (such as grooming). |
| Pacing | The individual continuously moves back and forth (such as jumping or traveling) between an area with a fixed pattern. |
| Panting | Open mouth respiration characterized by moderate to rapid airflow, |
| Shaking | The body of an individual trembles or shakes in place. |
| Yawning | The individual involuntarily opens their mouth, maximally widening the jaw, and takes in a deep inhale. This is followed by a sharp exhale. |

The RA’s noted the date of the video recording and the anxiety behavior displayed. Also, the duration of anxiety behaviors with a definitive start and end time were noted (sec.), these included: auto-grooming, auto-scratching, and examining genitalia. RA’s entered their observations on a Google Form, and the data were viewed on the resulting Google Spreadsheet. Each video file was observed by a minimum of two RA’s, and a data reliability check was conducted. Any video-file for which the two RA’s did not record the same data, were observed again by an additional RA or until the data recorded by two people was the same. Videos which did not include the focal individual, Medusa, or contained only periods of sleep, were excluded from analysis, as no relevant behaviors could be observed during these times.

### Statistical Analysis

Frequencies of each anxiety-related behavior were tallied across the three observation days. Because the data did not meet the assumptions of normality nonparametric statistics were used for the analysis. However, the common assumptions of nonparametric tests are independence and randomness, which were not met with this data because of the small sample size (n=1). Therefore, the results should be interpreted as within-individual behavioral variation and not a population-level pattern.

Pearson’s Chi-square tests were used to determine whether the frequency of anxiety-related behaviors differed within each day and across across three days peripartum (June 6 - June 7, 2017). Expected values were calculated under the assumption that all behaviors occurred with equal probability relative to the total number of bahviors observed per day. To control for multiple comparisons, the significance threshold was adjusted using Bonferroni’s correction (α = 0.05/11behaviors ≈ 0.0045).

Pairwise Chi square tests with Bonferroni adjustment for multi-comparisons (α = 0.05/3 comparisons ≈ 0.0167) were then conducted to compare behavioral frequencies between days (June 6 vs June 7; June 6 vs June 8; June 7 vs June 8) to determine which behaviors were performed significantly more on any given day. Expected frequencies for these pairwise tests were calculated under the assumption that each behavior would occur in equal proportion (in regard to total behaviors observed that day) across the two days being compared. When Chi-square test assumptions were violated due to low expected frequencies (< 5), two-tailed Fisher’s Exact Tests were applied instead. If the statistical tests resulted in significance, the difference between observed and expected counts (O-E) was used to identify the day on which each behavior occurred more frequently.

Fisher’s Exact Test was used to assess whether the frequency of anxiety-related behaviors was dependent on the day they occurred during the peripartum period. Because the contingency table was large (3 days x 11 behaviors), exact computation was not feasible and therefore the p-values were approximated using the Monte Carlo simulation with 1,000,000 replicates.

In addition, the duration of auto-grooming, auto-scratching, and examining genitalia were analyzed using the Kruskal–Wallis test to evaluate whether the distribution of bout durations differed across days. When significance was detected, pairwise Mann-Whitney U tests (equivalent to the Wilcoxon Rank-Sum Test) were conducted to determine which days differed significantly in duration.

## Results

### Frequency of maternal anxiety-related behaviors peripartum

The mother aye-aye displayed a total of 793 anxiety behaviors peripartum (**Figure 2**). The most observed behaviors were constructing and deconstructing nests (n=238), auto-scratching (n=229), and auto-grooming (n=131). The mother was also hyper-vigilant (n=97) and frequently examined genitalia (n=68). Less frequntly observed behaviors were nudging infant (n=14), pacing (n=7), shaking (n=6), and yawning (n=3). Panting and hair picking were not observed and were thus omitted from statistical analysis (**Figure 2**).

**Figure 2.**
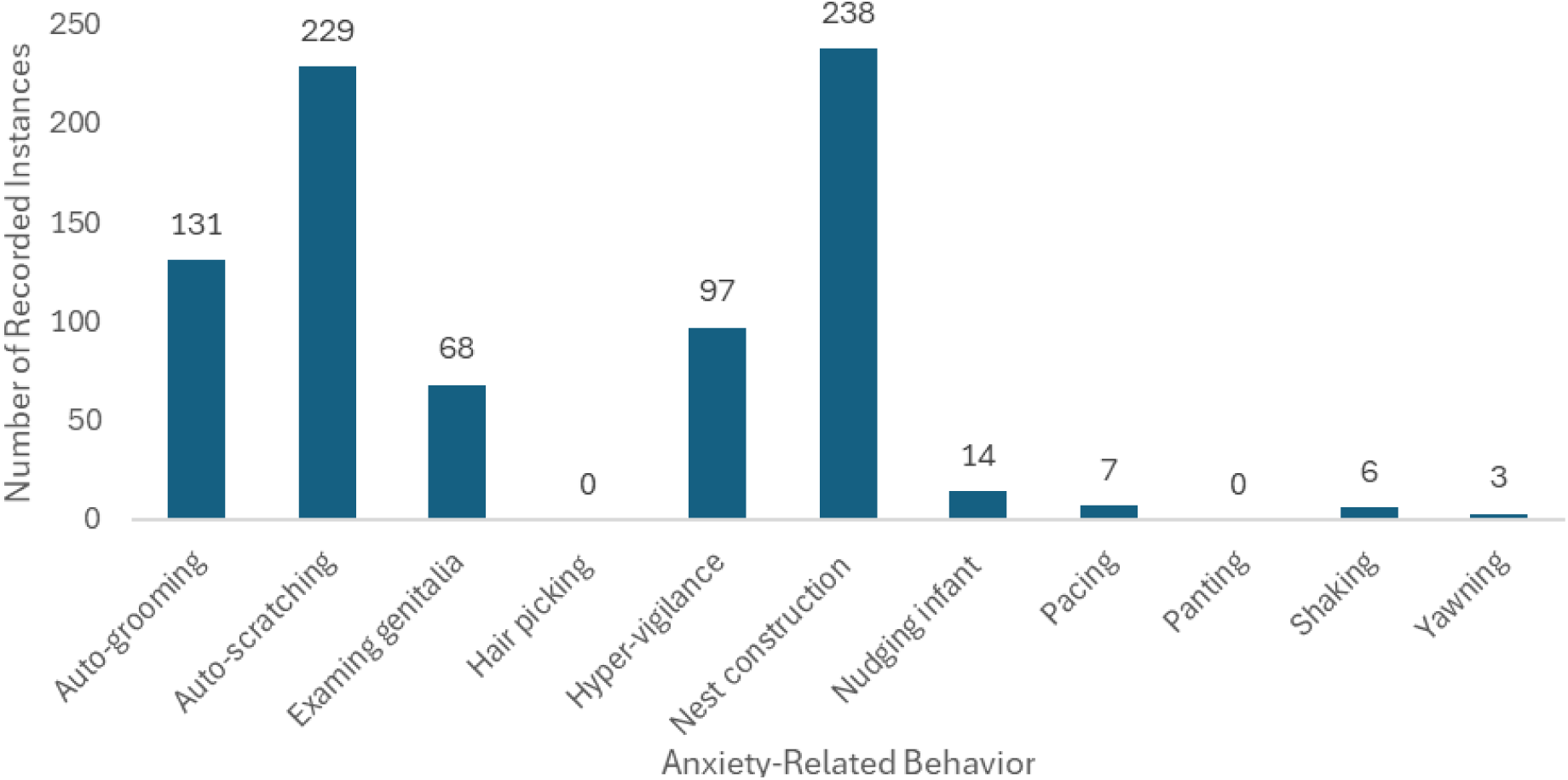
Total frequency of anxiety-related behaviors across the 3-day peripartum period in a mother aye-aye. Total recorded instances of 11 anxiety-related behaviors observed in the mother aye-aye, *Medusa*, across three consecutive peripartum days (June 6–8, 2017). Each bar represents the sum of behaviors recorded during 24 hours of continuous video footage per day (n = 1 subject, over 3 days), illustrating shifts in maternal anxiety-related behavioral expression across the peripartum period.

On the day before birth (June 6), the mother aye-aye engaged in eight types of anxiety behaviors totaling 484 occurrences and the frequency of these behaviors were significantly different (χ² = 1100.05, df = 10, p < 0.001). The mother engaged in the following: nest construction (n=212x, 43.8%), auto-scratching (n=136x, 28.1%), auto-grooming (n=57x, 11.8%), hyper-vigilance (n=52x, 10.7%), examining genitalia (n=16x, 3.31%), shaking (n=6x, 1.24%), yawning (n=3x, 0.62%), and pacing (n=2x, 0.41%) (**Figure 3a, Table S1**). On the day of birth (June 7), the mother aye-aye was seen performing five anxiety behaviors with 217 recorded instances which were also significantly different (χ² = 496.01, df = 10, p < 0.001). The mother participated in the following: auto-scratching (n=87x, 40.1%), auto-grooming (n=60x, 27.7%), examining genitalia (n=52x, 24%), hyper-vigilance (n=13x, 6%), and pacing (n=5x, 2.3%) (**Figure 3b, Table S1**). The mother aye-aye engaged in five anxiety behaviors which were significantly different (χ² = 162.51, df = 10, p < 0.001) on the day after birth (June 8) with a total of 92 recorded instances. These were hyper-vigilance (n=32x, 34.8%), nest construction (n=26x, 28.3%), nudging infant (n=14x, 15.2%), auto-grooming (n=14x, 15.2%), and auto-scratching (n=6x, 6.52%) (**Figure 3c, Table S1**).

**Figure 3:**
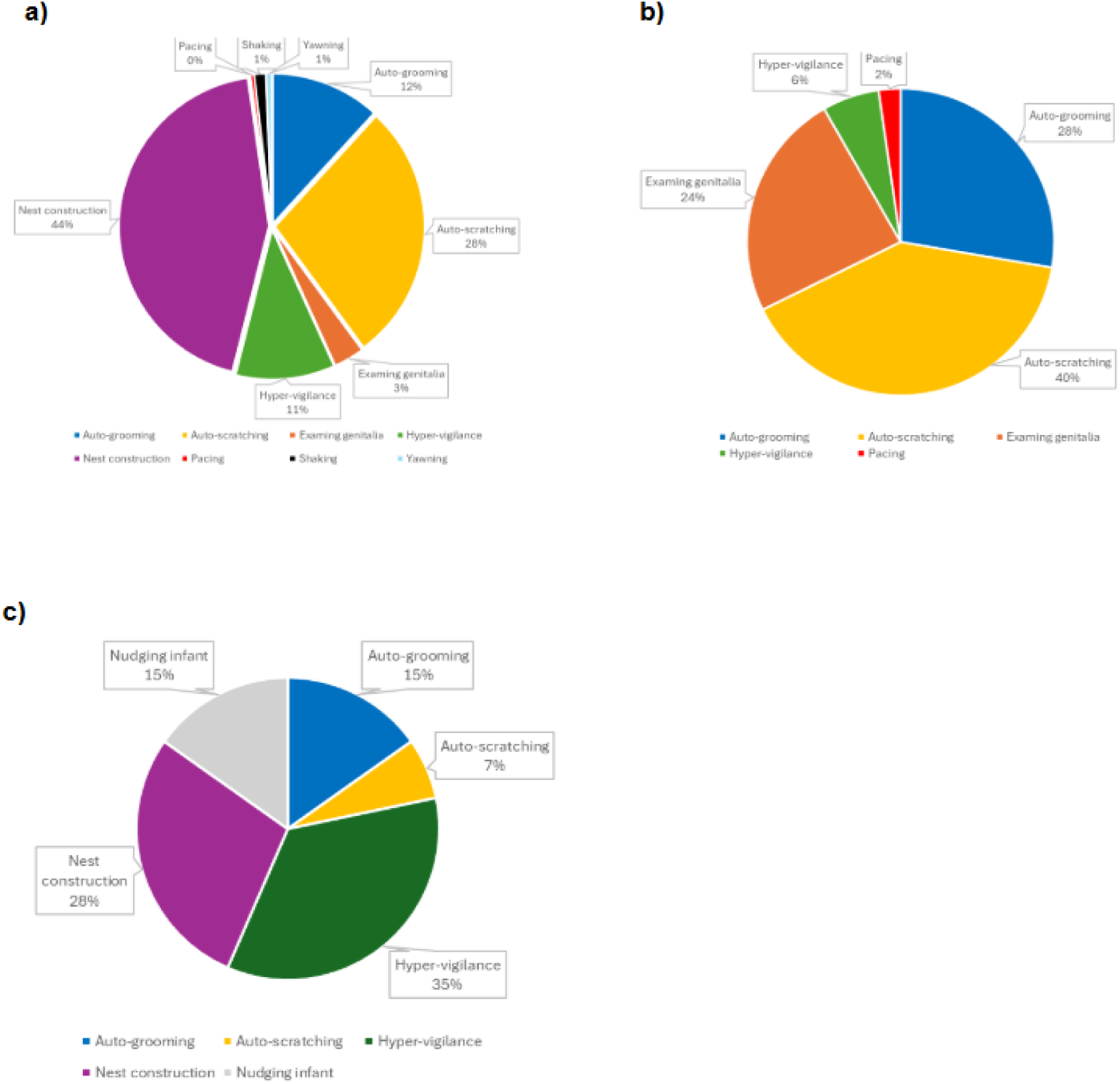
Percentage of maternal anxiety-related behaviors recorded on a given day peripartum (June 6 - June 8, 2017) for a mother aye-aye. Pie charts display percentage of time the mother aye-aye (n=1) engages in the anxious behavior peripartum. **a)** Day before birth, June 6 (χ² = 1100.05, df = 10, p < 0.001) **b)** Day of Birth, June 7 (χ² = 496.01, df = 10, p < 0.001) **c)** Day after birth, June 8 (χ² = 162.51, df = 10, p < 0.001). Percentages were calculated from the total instance of anxiety-related behaviors performed on a given day (24 hours of continuous video footage) and significance was corrected with Bonferroni adjustment (α =0.0045). See Table S1 for statistical details.

Chi-square goodness of fit tests (adjusted α = 0.0167) which were used to determine if the frequency of a behavior differed across three days also showed that auto-grooming (χ² = 30.34, df = 2, p < 0.001) and auto-scratching (χ² = 112.93, df = 2, p < 0.001) were statistically different (**Figure 4, Table S2**). Also, examining genitalia (χ² = 62.59, df = 2, p < 0.001), hyper-vigilance (χ² = 23.53, df = 2, p < 0.001), nest construction (χ² = 337.04, df = 2, p < 0.001), nudging infant (χ² = 28, df = 2, p < 0.001), and shaking (χ² = 12, df = 2, p < 0.001) were significantly different (**Figure 4, Table S2**). The frequency of pacing (χ² = 5.43, df = 2, p = 0.066) and yawning (χ² = 6, df = 2, p = 0.050) were not significantly different across the three-day study period (**Figure 4, Table S2**).

**Figure 4.**
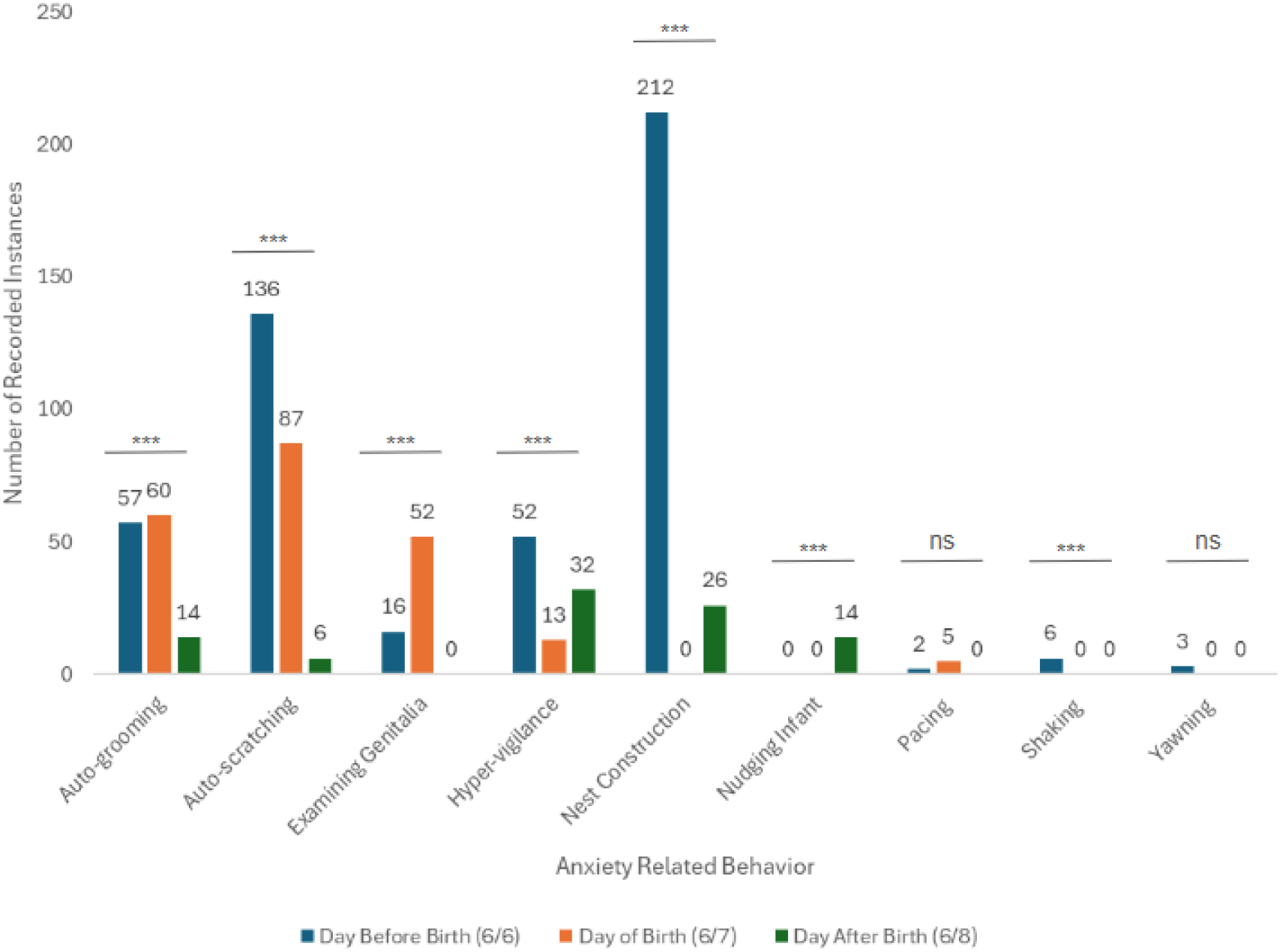
Frequency of nine maternal anxiety-related behaviors peripartum in a mother aye-aye. Number of recorded instances of nine maternal anxiety-related behaviors: self-grooming, self-scratching, examining genitalia, hyper-vigilance, nest construction, nudging infant, pacing, shaking, and yawning, on the day before (June 6), the day of (June 7), and the day after parturition (June 8). Each bar represents the total frequency of the behavior observed on the given day during 24 hours of continuous video footage per day (n = 1) reflecting day-to-day variation in total behavior. Statistical significance is indicated as **P < 0.01, ***P < 0.001, ns = not significant. Results are shown with Bonferroni adjustment for multiple comparisons (α = 0.0167). See Table S2 for statistical details.

### Pattern of maternal anxiety behavior peripartum

Post Hoc pairwise Chi-square tests with Bonferroni-adjusteed significance levels (α = 0.0167) were conducted to identify which days differed significantly in anxiety-related behavioral frequencies. Auto-grooming significantly differed between June 6 vs June 7 (χ² = 27.15, p < 0.001) and approached statistical significance between June 7 vs June 8 (χ² = 5.48, p = 0.019) (**Figure 5a, Table S3**). Comparison of observed and expected counts indicated that auto-grooming was significantly more frequent on the day of birth (June 7; p < 0.001). Auto-scratching was significantly different between all three days: June 6 vs June 7 (χ² = 9.93, p = 0.0016), June 6 vs June 8 (χ² = 19.38, p < 0.001), and June 7 vs June 8 (χ² = 34.61, p < 0.001) (**Figure 5b, Table S3**). Auto-scratching was significantly (p < 0.001) more frequent on the day of birth (June 7; p < 0.001), occurring at the highest proportional rate on this day. Examining genitalia was significantly different between June 6 vs June 7 (χ² = 161.76, p < 0.001) and June 7 vs June 8 (χ² = 51.34, p < p < 0.001) and was most frequent the day of birth (June 7; p < 0.001) (**Figure 5c, Table S3**). Hyper-vigilance was only significantly different between June 6 vs June 8 (χ² = 35.86, p < 0.001) and June 7 vs June 8 (χ² = 34.05, p < 0.001) (**Figure 5d, Table S3**).

**Figure 5.**
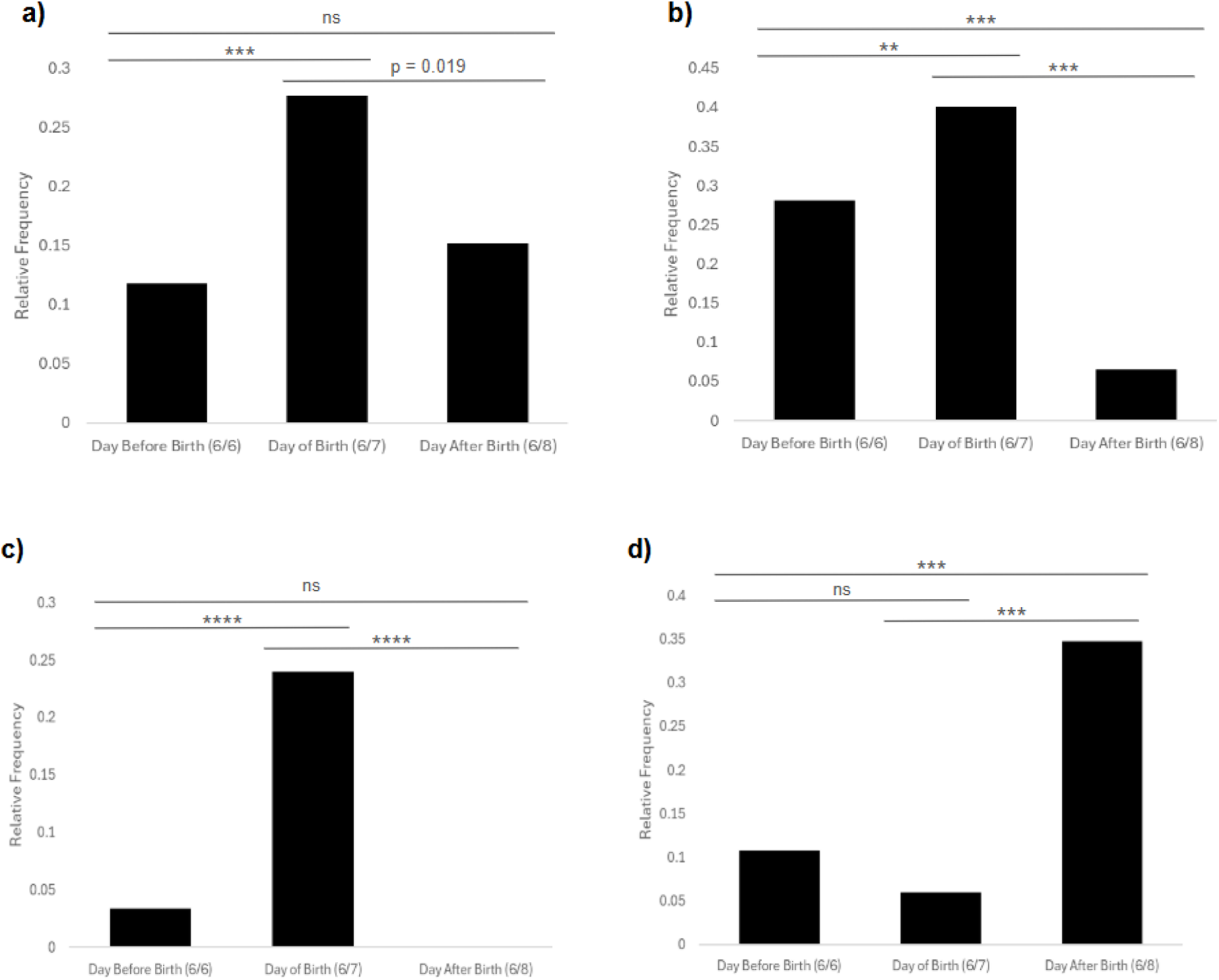
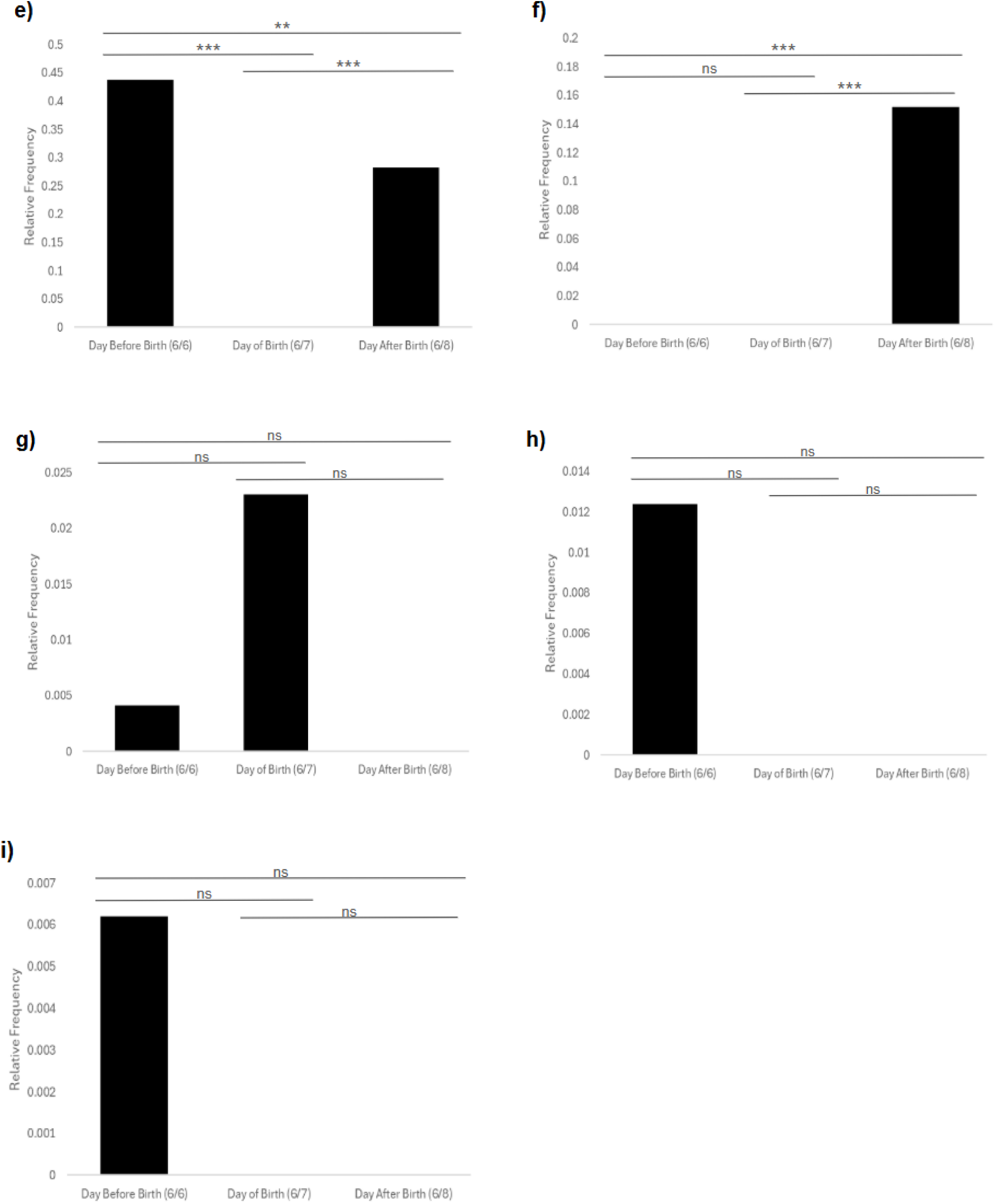
Relative frequency of maternal anxiety-related behavior peripartum for a mother aye-aye. Proportion of each behavior shown on the day before (June 6), the day of (June 7), and the day after parturition (June 8) for a mother aye-aye (n=1). Behaviors include **a)** self-grooming, **b)** self-scratching, **c)** examining genitalia, **d)** hyper-vigilance, **e)** nest construction, **f)** nudging infant, **g)** pacing, **h)** shaking, and **i)** yawning. Each bar indicates the relative frequency of the behavior across three observation days with 24 hours of continuous video-footage. Proportions were calculated as the number of instances divided by the total number of all behaviors recorded that day. This reflects the pattern of anxiety-related behavior the mother displayed, highlighting day-to-day differences. Data was collected with 24 hours of continuous video footage per day (n = 1 subject, over 3 days). Statistical significance is indicated as **P < 0.01, ***P < 0.001, ns = not significant. Results are shown with Bonferroni adjustment for multiple comparisons (α = 0.0167). See Table S4 for statistical details

Proportionally, hyper-vigilance was significantly most frequent the day after birth (June 8; p < 0.001). Finally, nest construction significantly differed between all days; June 6 vs June 7 (χ² = 136.26, p < 0.001), June 6 vs June 8 (χ² = 7.70, p = 0.006), and June 7 vs June 8 (χ² = 66.96, p < 0.001) (**Figure 5e, Table S3**) and was most frequent the day before birth (June 6; p < 0.01). Fisher’s exact tests further determined that nudging infant significantly differ between June 6 vs June 8 (p < 0.001) as well as June 7 vs June 8 (p < 0.001) (**Figure 5f, Table S4**). Examining genitalia was only seen the day after birth, thus most frequent on this day (June 8; p < 0.001). The remaining behaviors: pacing, shaking, and yawning did not significantly differ between days (p > 0.0167) (**Figure 5g - 4i, Table S4**).

Fisher’s exact test, approximated using Monte Carlo simulation with 1,000,000 replicates, indicated that the distribution of behaviors was related to the day they happened (p < 0.001). This result supports the alternative hypothesis that the behavioral frequencies depended on the day of observation peripartum rather than random variation. As a two-sided test, it suggests that at least one behavior occurred at a significantly different relative frequency between days.

### Duration of maternal anxiety-related behavior peripartum

The mean duration of auto-grooming was 16.49 seconds over the three-day study period and auto-scratching and examining genitalia were 9.62 and 13.74 respectively. The distribution of bout duration also changed throughout the study period (**Figure 6**). On the day before birth, the average duration of auto-grooming, auto-scratching, and examining genitalia was 24.09 seconds, 11.35 seconds, and 13.5 seconds, respectively (**Figure 6a**). On the day of birth, the average duration of auto-grooming, auto-scratching, and examining genitalia was 26.03 seconds, 8.71 seconds, and 20.13 seconds respectively (**Figure 6b**). On the day after birth, the average duration of auto-grooming, auto-scratching, and examining genitalia was 26.57 seconds, 11 seconds, and 0 seconds respectively (**Figure 6c**). The Kruskal Wallis test showed that there was a significant difference in the duration of auto-scratching (p = 0.032, **Table S5**), however post hoc analysis showed that no two days were significantly different pairwise (p > 0.05).

**Figure 6.**
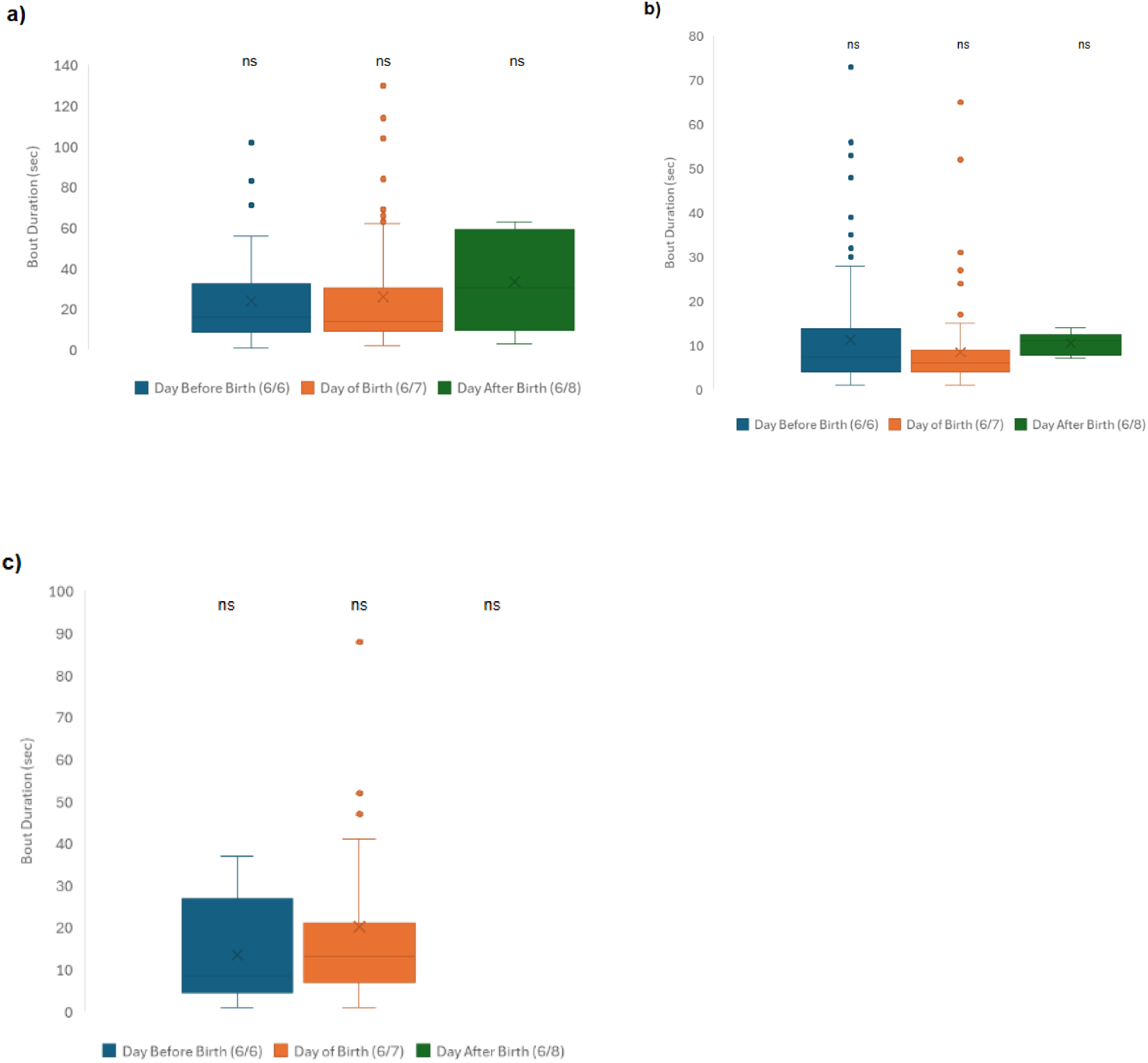
Duration of auto-grooming, auto-scratching, and examining genitalia peripartum for a mother aye-aye. Box and whisker plots show that distribution of anxiety-related behaviors on **a)** the day before birth (June 6) **b)** the day of birth (June 7), and **c)** the day after birth (June 8) for the mother aye-aye (n=1, 24 hour video-footage). ns = nonsignificant. See Table S5 for statistical details.

Auto-grooming (p = 0.262, **Table S5**), and examining genitalia (p = 0.285, **Table S5**) durations across the three days peripartum were also not significant. This indicates that the duration of auto-grooming, auto-scratching, and examining genitalia remained mostly stable over time.

## Discussion

The mother engaged in high levels of anxiety-related behaviors peripartum. It is challenging, though, to determine whether exhibiting 793 anxiety behaviors peripartum is normal since similar data in nonhuman primates is lacking. However, previous research has demonstrated that maternal behavior can serve as an important indicator of emotional state in nonhuman primate mothers (Troisi et al., 1991; Maestripieri, 1993a; Marais et al., 2006; Bass, 2016). Using this framework, the behavioral patterns observed in an aye-aye mother at the Duke Lemur Center are interpreted as potential indicators of maternal anxiety.

### 1. Key Maternal Anxiety-Related Behaviors Peripartum

#### 1a. Nest-Construction

The mother engaged in constructing and deconstructing nests more than any other behavior. Nesting plays a key role in Malagasy primate life histories and has been associated with faster fetal and postnatal growth rates, as well as shorter interbirth intervals (Tecot et al., 2012). Although data on *Daubentonia* was scarce and therefore, not included in Tecot et al. ’s (2012) study, the strong selective pressures shaping nest-building in other strepsirrhines suggest that mothers frequently manipulate their nest as both an adaptive and stress-regulatory function. Thus, the repetitive nest construction observed in Medusa could reflect heightened arousal and/or an anxiety-related need to control the environment during the peripartum period.

#### 1b. Auto-Scratching

Auto-scratching was the next most common behavior the mother exhibited. This makes sense as it has long been established that scratching increases during stressful situations (e.g., Castles et al., 1999; Maestripieri et al., 1993a) and is one of the most commonly observed anxiety behaviors in nonhuman primates (Maestripieri et al., 1992). Among seven group-living macaque mothers, maternal scratching correlated with attentive and possessive maternal styles (Troisi et al., 1991). Further, in a study on ten group-living macaques, mothers who were physically abusive towards their infants during the first 2–3 weeks of life, failed to show the typical separation-related increase in scratching (Maestripieri, 1998). The high frequency of scratching observed here is therefore consistent with an anxious yet attentive maternal style and is a part of being a “good” mother.

#### 1c. Auto-grooming

Auto-grooming was also highly exhibited in the aye-aye mother. While this behavior is commonly characterized as hygienic, serving to remove dirt, ectoparasites, and defoliated skin (Struhsaker, 1967; Tanaka & Takefushi, 1993; Ventura et al., 2005), auto-grooming may also play a role in stress regulation across taxa. In socially stressed rodents, for instance, mothers neglected pup care in favor of self-grooming (Carini et al., 2013), while primates showed increased auto-grooming following infant separations or environmental disturbances (Klopfer & Dugard, 1976; Feistner & Ashbourne, 1994). Moreover, grooming behaviors in primates are physiologically associated with decreased stress hormone (glucocorticoid) levels (Gust et al., 1993) and increased production of endogenous opioids (β-endorphins) (Keverne et al., 1989). Together, these findings suggest that elevated auto-grooming in the aye-aye mother is a likely important indicator of maternal anxiety.

#### 1d. Hyper-vigilance

Frequent hyper-vigilance suggests that the mother maintains a heightened awareness of her surroundings, consistent with an adaptive anxiety response. Heightened vigilance is advantageous in contexts involving potential threats (Gaynor & Cords, 2012; MacIntosh & Sicotte, 2009), and is particularly evident in mothers monitoring infants (Kutsukake, 2006; Maestripieri, 1993b). Remaining motionless and alert can prepare the individual to respond quickly to danger and may reduce the likelihood of detection by predators (Coleman & Pierre, 2014). For example, free-ranging golden snub-nosed monkey (*Rhinopithecus roxellana)* mothers exposed to traumatic events such as infanticide showed increased hyper-vigilant and protective behaviors (Zhao et al., 2025). Such behaviors can also shape offspring threat sensitivity later in life (Mandalaywala et al., 2014). Collectively, maternal vigilance appears to serve an adaptive role, enhancing offspring protection and shaping developmental responses to threat. This may help explain why Medusa frequently engaged in hyper-vigilant behaviors during peripartum.

#### 1e. Examining Genitalia

Examining genitalia was another behavior exhibited by the anxious mother aye-aye. Self-examination of the genital area, observed primarily around parturition, is documented in several primate species and typically reflects a normal component of the birthing process (Solanki & Zothansiama, 2013; Turner et al., 2010; Nguyen et al., 2017). In captive stump-tailed macaques (*M. arctoides*), throughout labor and delivery, mothers were observed repeatedly licking and inspecting their genital region (Solanki & Zothansiama, 2013). While self-touching behaviors can be associated with anxiety (Schino et al., 1996; Troisi, 2002), the pattern observed here more likely represents parturition-related self-assessment and hygiene rather than an anxiety-specific behavior.

### 2. Duration of Anxiety-Related Behaviors

In addition to the frequency of anxiety-related behaviors, the mean duration of behaviors such as auto-grooming, auto-scratching, and examining genitalia could provide valuable insights into maternal anxiety. Of the three behaviors examined, only the mean duration of auto-scratching differed significantly across the peripartum period. Scratching duration decreased on the day of birth and then increased slightly the following day, suggesting fluctuations in the intensity or persistence of maternal anxiety. Because scratching is a well-established indicator of arousal in nonhuman primates (Aureli & van Schaik, 1991; Maestripieri et al., 1992; Troisi, 2002), variation in its mean duration may reflect differences in how long the mother remained in an anxious state after engaging in the behavior. Longer durations of anxiety-related behaviors may be interpreted as reflecting heightened arousal whereas shorter durations may indicate momentary reactivity or rapid recovery and habituation (Aureli, 1992; Diezinger & Anderson, 1986; Aureli & van Schaik, 1991).

Non-significant changes in the mean duration of auto-grooming and examining genitalia may suggest that these behaviors are less sensitive to the stimuli present peripartum and/or that the behaviors may have some physical constraint resulting in the inability to continue the behavior for longer durations (e.g., presence of infant, limited mobility inside the nest). However, the short observation period and small sample size may further limit interpretation. Additional research is needed to determine how behavioral duration, alongside frequency, can serve as an index of maternal anxiety.

### 3. Behavioral Patterns of Anxiety Peripartum

The mother aye-aye engaged in high levels of anxiety-related behaviors during the peripartum period, displaying distinct behavioral patterns across days. These behaviors, such as nesting, auto-scratching, auto-grooming, hyper-vigilance, and genital examination, likely reflect context-dependent responses to the stressors associated with parturition and infant care. Other studies have shown that maternal anxiety behaviors are context-dependent and mothers may adjust their behavior given the stimuli they face (Maestripieri, 2011; Stanton et al., 2015). Free ranging Tibetan macaque mothers (*Macaca thibetana*), for example, change their visual monitoring and protective behaviors when the perceived social risk to their infant is high (Liu et al., 2024) and wild chimpanzee (*P. troglodytes*) mothers are more vigilant during infant separation events (Kutsukake, 2006). Other behaviors like examining the genitalia also seem to be associated with the stressor present. In Japanese macaques *(M. fuscata*) and long-tailed macaques (*M. fascicularis*) mothers examine genitalia more frequently and more intensely around the time of parturition (Turner et al., 2010; Kemps & Timmermans, 1982). Likewise, maternal physiological stress (fecal/glucocorticoid measures) covaries with day-to-day maternal behaviour in wild populations of chimpanzees (*P. troglodytes*), indicating that maternal state varies with current stressors (Stanton et al., 2015).

Daily variation in the aye-aye mother’s behavior followed this pattern, as maternal state was dependent on the observation day. These findings indicate that maternal anxiety behaviors are influenced by the stimuli and physiological stressors present. This may also explain the absence of certain behaviors, such as genital examination, following the resolution of the stressors.

#### 3a. Day Before Birth (June 6): Nest Construction and Preparation

On the day before birth (June 6, 2017) where the mother spent most of her time constructing and deconstructing the nest (43.8%). This could indicate that the mother is preparing for parturition and creating a safe and comfortable environment for herself and the infant. Nesting provides security and comfort and can reduce stress before birth (Stewart et al., 2007; Fruth et al., 2018). In the wild, safe nesting sites are essential for protecting vulnerable infants from predators (Stewart & Pruetz, 2013). Auto-scratching also occurred frequently (28.1%), consistent with its role as a common anxiety-related behavior in primates (Maestripieri, et al., 1992). The combination of heightened nesting and scratching likely reflects a mix of anticipatory arousal and environmental preparation preceding birth.

#### 3b. Day of Birth (June 7): Hygiene and Parturition Behaviors

On the day of birth (June 7, 2017) the mother aye-aye spends most of her time auto-scratching (40.1%) and auto-grooming (27.6%). Auto-scratching has been found to be a useful tool to investigate maternal behavior in nonhuman primates and this behavior often becomes more frequent when mothers sense threats and/or are anxious over their infant (Maestripieri, 1993a; Troisi et al., 1991). Both behaviors serve hygienic and stress-regulatory functions, aiding in scent eradication, cleanliness, and potentially predator avoidance (Feusner et al., 2009; Spruijt et al., 1992). Grooming after birth is particularly important for reducing infection risk for both mother and infant (Akinyi et al., 2013; Maródi, 2006) and may also provide nutritional or physiological benefits through ingestion of birthing fluids and placenta (Fujisawa et al., 2016; Bridges, 2015). The mother also frequently examined her genital region (24.0%), a behavior common among primate mothers around the time of parturition. This behavior is often associated with the discomfort of birth, swelling and irritation of the genitals, and nutrient retrieval from birthing fluids (Kemps & Timmermans, 1982; Tinklepaugh & Hartman, 1930). Other primate mothers focus their attention on the genital area around the time of birth to assess the infant’s position, physical changes of the birthing process, and to increase the flexibility of the tissues for birth (macaques: Solanki & Zothansiama, 2013; Turner et al., 2010; Kemps & Timmermans, 1982; bonobos: Demuru et al., 2018; geladas: Nguyen et al., 2017). Because genital examination is widely observed in nonhuman primates around parturition and has not been characterized as anxiety-related or stereotypic, it likely represents a normal component of the birthing process rather than an anxiety-related behavior.

#### 3c. Day After Birth (June 8): Hypervigilance and Maternal Protection

Finally, on the day after birth (June 8, 2017) the mother spends most of her time hyper-vigilant (34.8%). This suggests a transition to maternal protectiveness and alertness over the infant. Heightened vigilance after parturition may facilitate early recognition of threats to the newborn (Maestripieri, 1993b; Kimble et al., 2013) and has been observed in primate mothers when predation or social risks are high (Liu et al., 2024; Kinally et al., 2019). This pattern indicates a shift in maternal anxiety behaviors from self-directed behaviors (e.g., scratching and grooming) to alertness over the new infant. Vigilance may thus indicate an anxious mother.

It is important to note that the behavioral data for this day may underestimate activity due to methodological constraints. Following birth, the mother was placed in a large carrier on the enclosure floor. The carrier substantially limited visibility of both the mother and her infant, obstructing the coder’s view and potentially leading to missed or inaccurately coded behaviors. This reduced visibility likely contributed to the reduced number of behaviors recorded (92 compared to 484 and 217 on the previous days). It is possible that some behaviors may have occurred out of the camera’s field of view..

### 4. Adaptive Hypothesis of Maternal Anxiety

The observed behavioral patterns support the adaptive hypothesis of maternal anxiety, which proposes that moderate levels of maternal anxiety enhance offspring survival by promoting vigilance, responsiveness, and attentiveness. Maternal anxiety can increase responsiveness to threats (Bar-Haim et al., 2007) and strengthen affiliative behaviors such as attending to infant distress signals (Nguyen et al., 2008). In yellow baboons (*Papio cynocephalus*), females with elevated prepartum glucocorticoid levels showed greater responsiveness to infant distress calls than those with lower levels (Nguyen et al., 2008). Similarly, mothers with higher prepartum cortisol displayed increased affiliative behaviors with infants, whereas elevated postpartum cortisol was associated with reduced attentiveness (Bardi et al., 2004).

Experimental evidence from rodents also demonstrates adaptive effects of maternal stress on offspring. For example, in red squirrels (*Tamiasciurus hudsonicus*), mothers with higher glucocorticoid levels due to population density stress produced faster-growing offspring, consistent with the adaptive maternal effect hypothesis (Dantzer et al., 2013). Across taxa, maternal anxiety and stress can influence offspring physiology and behavior in ways that improve coping with environmental challenges (Love et al., 2013; Bauer et al., 2019; Gluckman & Hanson, 2004; Barker, 2002; Lee & Zucker, 1988). In our present study, the aye-aye mother’s behavioral pattern aligns with this adaptive framework. The day before birth, the mother prepares her environment, the day of birth, she displays heightened arousal for parturition and increased hygiene and self-directed behaviors, and the day after birth, the mother is vigilant of her infant. This suggests that maternal anxiety may function to promote both mother and infant protection and survival.

### 5. Behaviors Not Associated with Maternal Anxiety

The mother did not engage in panting or hair picking and infrequently engaged in nudging the infant, pacing, shaking, or yawning. This suggests that these behaviors are not characteristic of maternal anxiety in aye-ayes.

#### 5a. Absent Behaviors: Panting and Hair Picking

Panting was absent and is rarely observed in primates outside thermoregulatory functions (Hildwein & Goffart, 1975; Maho et al., 1981). Panting as an anxious behavior is more commonly seen in canines especially those with noise sensitivity and separation events (Tiira et al., 2016; Overall et al., 1997). Also, although hair picking occurs in anxious nonhuman primates, particularly in females (Reinhardt, 2005), there are few studies linking hair picking to maternal anxiety. Hair picking is more closely linked to the stress of captivity, poor welfare, and social environment (Brand & Marchant, 2015; Heagerty et al., 2017). Considering that zero instances of panting and hair picking were observed in the mother, it is unlikely that these behaviors indicate maternal anxiety in aye-ayes.

#### 5b. Infrequent Behaviors: Nudging Infant, Pacing, Shaking, and Yawing

Of the behaviors that happened infrequently, nudging or repositioning the infant likely represents normal maternal caregiving rather than anxiety. Mothers often interact, contact and reposition their infant, doing so as a form of maternal caregiving (Hansen, 1966; Maestripieri et al., 2008), protecting the infant (Hansen, 1966; Schino et al., 2003; Maestripieri, 2001), and encouraging locomotion in their infants (Maestripieri, 1995, 1996). Thus, Medusa was likely nudging her infant (infrequently) to reposition her and not because Medusa was particularly anxious at that moment.

Pacing was rarely observed, and the pacing bouts that were observed may represent short episodes of *agitated locomotion* rather than stereotypic pacing. Poirier et al. (2019) emphasized that pacing involves long (up to 10 min.) invariant routes, whereas agitated locomotion is shorter (< 20 sec) with a varied path. Also, Poirier et al. (2019) found that pacing did not increase during or after agonistic interactions, but agitated locomotion did increase during agonistic interactions. These results suggest that agitated locomotion is an indicator of acute stress. Medusa seemed to engage in agitated locomotion (albeir rare), rather than pacing. Future research should therefore adopt the stricter definition of pacing outlined by Poirier et al. (2019) and consider both enclosure constraints and contextual stressors when interpreting locomotor behaviors.

Many mammals such as dogs (Grigg et al., 2021) and nonhuman primates (Novak et al., 2014) shake when anxious. Shaking is the body’s response that releases stress hormones and tension. Medusa did not shake frequently, indicating that her sympathetic nervous system and stress circuits were likely not activated.

Medusa was rarely observed yawning, which is a common physiological response to stress and anxiety, and helps the body self-regulate. In primates, yawning is context-dependent and is often reflective of agonistic tension and arousal from unfamiliar events (Vick et al., 2010; Leone et al., 2014; Palagi et al., 2009; Paukner & Anderson, 2006). Yawning is also commonly correlated with circadian fluctuations (Landolt et al., 1995; Depute, 1994; Vick & Paukner, 2010). In humans, yawning is more contagious in pregnant women than nulliparous women (Norscia et al., 2021). So, the fact that Medusa was not observed yawning a lot may not be that unusual as it could indicate sleep-wake cycles or context-specific changes related to her maternal state.

#### 5c. Observational Limitations

The lack of these behaviors may also indicate some human error. Because aye-aye’s are nocturnal and live in cocoon-like nests, it is often difficult to observe them properly even with the use of video footage. Some behaviors could have been easily overlooked or obstructed from view and therefore not recorded.

### 6. Limitations

Several limitations in addition to the environmental difficulties of a nocturnal, nest dwelling primate, should be considered when interpreting these findings. First, human error may have influenced data collection, as observational studies are inherently subject to bias. Behavioral definitions can vary between coders; however, each video file was coded at least twice, and coder comparisons were conducted to minimize discrepancies. When disagreement persisted, an experienced coder rewatched the footage until consensus was reached.

Second, the study covered only three days surrounding parturition, providing a limited view of the mother’s behavioral repertoire. To fully understand the temporal dynamics of maternal anxiety, future work should include continuous 24-hour observations both before and several months after birth. Prepartum observations will establish a baseline and postpartum data would help determine whether anxiety-related behaviors are uniquely elevated around the time of birth or part of a broader pattern of maternal behavior.

Third, the captive environment may have influenced the expression and frequency of anxiety-related behaviors. Captivity can induce abnormal or constrained behaviors due to environmental stressors and limited space (Nowak et al., 2000). External stressors unrelated to birth may therefore have contributed to the observed behavioral patterns. In addition, Duke Lemur Center husbandry staff were seen entering the enclosure on many occasions to check the wellbeing of the mother and her infant which could have exacerbated anxiety behaviors. Future research should attempt to disentangle anxiety associated with parturition from anxiety resulting from captivity.

Finally, the study’s small sample size (n = 1) prevents generalization to the wider aye-aye population and limits assessment of individual variation among mothers. Despite this, the present work provides the first detailed documentation of maternal anxiety in this species and offers a foundation for future comparative studies.

### 7. Significance and Implications

This report is the first study of maternal anxiety in an aye-aye mother. Because research on anxiety in nonhuman primates, particularly in aye-ayes is so limited, this study contributes to a broader understanding of how maternal anxiety is expressed across primate species. Anxiety in non-human primates is difficult to define because it manifests in a wide array of behaviors and may coexist with other underlying issues (Coleman & Pierre, 2014). By characterizing the behavioral repertoire of an anxious aye-aye mother, this study helps identify which stressors elicit specific anxiety responses and provides valuable insight into how nonhuman primates display stress. Such knowledge can inform both scientific understanding and the development of more effective husbandry and welfare practices for captive populations. Understanding how aye-aye mothers respond to stressors is especially important in captive populations, where welfare is critical to the success of conservation programs for this endangered species (Louis et al., 2020)

Stress and anxiety are adaptive in moderation, helping individuals respond to environmental or social challenges and promoting vigilance and maternal attentiveness. However, when these states become chronic or repetitive, they may develop into stereotypies—behaviors that are invariant, functionally associated with coping, and maladaptive over time (Mason, 1991; Wielebnowski, 2003). Chronic stress can lead to “distress,” impairing the individual’s ability to recover from stimuli and causing physiological and behavioral dysfunction, including immune suppression and reproductive inhibition (Wielebnowski, 2003).

In this study, the anxiety-related behaviors observed appear to reflect short-term, parturition-dependent responses rather than chronic stereotypies. Further research incorporating baseline and postpartum data will be necessary to confirm whether these patterns persist or remain adaptive.

Finally, maternal stress can have far-reaching implications for both the mother and her offspring. Elevated maternal stress hormones have been associated with reduced maternal care, increased rejections, and negative effects on infant development and immune function (Bardi et al., 2004; Saltzman & Abbott, 2009; Coe et al., 2002; Reyes & Coe, 1997). Identifying the behavioral indicators of maternal anxiety in aye-ayes therefore not only advances scientific understanding of primate stress responses but also provides essential tools for improving captive management and promoting the wellbeing of both mothers and infants.

### 8. Future Directions

Future research should extend these findings by incorporating longer observation periods, including both prepartum and postpartum data, to assess whether anxiety-related behaviors persist or fluctuate over time. This information could help determine whether the observed behaviors represent transient, acute responses or develop into chronic patterns consistent with stereotypies. These future investigations should also distinguish between acute, adaptive anxiety responses and chronic, repetitive stereotypies, which may signal compromised welfare. Defining these thresholds of maladaptive behavior would allow husbandry staff, in the context of captivity, to respond more efficiently and appropriately to behavioral changes.

Additionally, expanding the dataset to include multiple individuals will allow for population-level comparisons and identification of individual variation in maternal anxiety. Incorporating physiological measures in future studies, such as glucocorticoid concentrations, would also provide valuable physiological measures of anxiety which can be compared with the behavioral output of the mother(s). This would contribute to insight of the hormonal correlates of behavioral anxiety indicators.

Future research should also investigate the factors that mitigate stress in captive aye-ayes. Studies across taxa demonstrate that both pharmacological and environmental interventions can reduce anxiety-related or stereotypic behaviors. For example, the administration of anxiolytic drugs such as lorazepam or diazepam has been shown to selectively diminish displacement activities, including scratching, particularly among low-ranking or socially stressed primates (Schino et al., 1991, 1996; Cilia & Piper, 1997; Barros et al., 2000). Environmental enrichment, such as opportunities for object manipulation, has likewise been found to alleviate anxiety in socially isolated primates (Berkson et al., 1963).

Applying these insights to aye-aye management could help identify welfare practices that reduce anxiety and prevent the emergence of stereotypies. Future work should therefore integrate behavioral, physiological, and environmental data to develop comprehensive strategies for supporting maternal wellbeing and infant development in captivity. These future directions are essential for linking behavioral indicators of anxiety with conservation-focused welfare management in captive populations.

## Conclusion

The aye-aye mother engaged in many anxiety behaviors peripartum which changed in frequency depending upon the stressor present that day. High levels of auto-scratching, auto-grooming, examining genitalia, nest construction, and hyper-vigilance may all indicate the mothers anxiety. On the day before birth, where she is preparing for parturition, there is an increase in nest construction. On the day of birth, here the mother is engaging in hygiene and nutrient retrieval behaviors, there is a high frequency of scratching, grooming, and examining genitalia. On the day after birth, where the mother is more alert of potential threats to her infant, she participates in high levels of hyper-vigilance. These anxiety-related behaviors may indicate an acute, adaptive response to stressors pertaining to motherhood. Understanding these behaviors will enable husbandry staff to better care for anxious mothers and prepare an environment that decreases the stress levels of expectant aye-aye mothers and prevent the initiation of more maladaptive stereotypies. This study also gives insight to the complex behavioral repertoire of anxious non-human primates.

## Acknowledgements

We would like to extend our gratitude to Lauren Boyd, Tiffany Brocco, Eli Benbenek, Hyde Parkinson, Kayla Ruff, Chloe Glynn, Thomas Sutton, Jonah Peckham, Ali Monahan, Sawyer Roy, and the many other research assistants who helped code countless hours of video footage and made this study possible. Special thanks to the Duke Lemur Center for their support. Funding and additional help was generously provided by NC State’s Provost’s Professional Experience Program (PEP), NCSU Office of Undergraduate Research (OUR), NCSU Department of Biological Sciences (DBS), and NCSU’s DBS’s Support for Undergraduate Research (SURE). Added thanks to the NCSU BIO 267/269 Research PackTrack Program for providing the space and foundation that allowed this project to begin.

## References

Akinyi, M. Y., Tung, J., Jeneby, M., Patel, N. B., Altmann, J., & Alberts, S. C. (2013). Role of grooming in reducing tick load in wild baboons (*Papio cynocephalus*). Animal Behaviour, 85(3), 559–568. 10.1016/j.anbehav.2012.12.012

Altmann, J. (1974). Observational study of behavior: Sampling methods. Behaviour, 49(3–4), 227–266. 10.1163/156853974X00534

Antelman, S. M., & Caggiula, A. R. (1980). Stress-induced behavior: Chemotherapy without drugs. In J. M. Davidson & R. J. Davidson (Eds.), The Psychobiology of Consciousness (pp. 65–104). Springer US. 10.1007/978-1-4684-3456-9_4

Araji, S., Griffin, A., Dixon, L., Spencer, S.-K., Peavie, C., & Wallace, K. (2020). An overview of maternal anxiety during pregnancy and the post-partum period. Journal of Mental Health & Clinical Psychology, 4(4). https://www.mentalhealthjournal.org/articles/an-overview-of-maternal-anxiety-during-pre gnancy-and-the-post-partum-period.html

Aureli, F. (1992). Post-conflict behaviour among wild long-tailed macaques (*Macaca fascicularis*). Behavioral Ecology and Sociobiology, 31(5). 10.1007/BF00177773

Aureli, F., Cords, M., & van Schaik, C. P. (2002). Conflict resolution following aggression in gregarious animals: A predictive framework. Animal Behaviour, 64(3), 325–343. 10.1006/anbe.2002.3071

Aureli, F., & Schaik, C. P. V. (1991). Post-conflict behaviour in long-tailed macaques (*Macaca fascicularis*): II. Coping with the uncertainty. Ethology, 89(2), 101–114. 10.1111/j.1439-0310.1991.tb00297.x

Aureli, F., Van Schaik, C. P., & Van Hooff, J. A. R. A. M. (1989). Functional aspects of reconciliation among captive long-tailed macaques (*Macaca fascicularis*). American Journal of Primatology, 19(1), 39–51. 10.1002/ajp.1350190105

Bailey, M. T., & Coe, C. L. (1999). Maternal separation disrupts the integrity of the intestinal microflora in infant rhesus monkeys. Developmental Psychobiology, 35(2), 146–155.

Baker, K. C., & Aureli, F. (1997). Behavioural indicators of anxiety: An empirical test in chimpanzees. Behaviour, 134(13–14), 1031–1050. 10.1163/156853997X00386

Ballard, C. G., Davis, R., Handy, S., & Mohan, R. N. (1993). Postpartum anxiety in mothers and fathers. The European Journal of Psychiatry, 7(2), 117–121.

Bardi, M., French, J. A., Ramirez, S. M., & Brent, L. (2004). The role of the endocrine system in baboon maternal behavior. Biological Psychiatry, 55(7), 724–732. 10.1016/j.biopsych.2004.01.002

Bar-Haim, Y., Lamy, D., Pergamin, L., Bakermans-Kranenburg, M. J., & van IJzendoorn, M. H. (2007). Threat-related attentional bias in anxious and nonanxious individuals: A meta-analytic study. Psychological Bulletin, 133(1), 1–24. 10.1037/0033-2909.133.1.1

Barker, D. J. P. (2002). Fetal programming of coronary heart disease. Trends in Endocrinology & Metabolism, 13(9), 364–368. 10.1016/S1043-2760(02)00689-6

Barnett, J. L., & Hemsworth, P. H. (1990). The validity of physiological and behavioural measures of animal welfare. Applied Animal Behaviour Science, 25(1), 177–187. 10.1016/0168-1591(90)90079-S

Barros, M., Boere, V., Huston, J. P., & Tomaz, C. (2000). Measuring fear and anxiety in the marmoset (*Callithrix penicillata*) with a novel predator confrontation model: Effects of diazepam. Behavioural Brain Research, 108(2), 205–211. 10.1016/S0166-4328(99)00153-9

Bass, A. (2016). *An Analysis of the Correlation between Cortisol Levels and Anxious Behavior of Captive Aye-Ayes (Daubentonia madagscariensis) at the Duke Lemur Center*. Honors thesis, Duke University. https://hdl.handle.net/10161/11984.

Bauer, C. M., Correa, L. A., Ebensperger, L. A., & Romero, L. M. (2019). Stress, sleep, and sex: A review of endocrinological research in *Octodon degus*. General and Comparative Endocrinology, 273, 11–19. 10.1016/j.ygcen.2018.03.014

Bayart, F., Hayashi, K. T., Faull, K. F., Barchas, J. D., & Levine, S. (1990). Influence of maternal proximity on behavioral and physiological responses to separation in infant rhesus monkeys (*Macaca mulatta*). Behavioral Neuroscience, 104(1), 98–107.

Berkson, G. (1967). Abnormal stereotyped motor acts. Proceedings of the Annual Meeting of the American Psychopathological Association, 55, 76–94.

Berkson, G. (1968). Development of abnormal setereotyped behaviors. Developmental Psychobiology, 1(2), 118–132. 10.1002/dev.420010210

Berkson, G., Mason, W. A., & Saxon, S. V. (1963). Situation and stimulus effects on stereotyped behaviors of chimpanzees. Journal of Comparative and Physiological Psychology, 56(4), 786–792. 10.1037/h0044086

Bhargava, D., & Trivedi, H. (2018). A study of causes of stress and stress management among youth. IRA-International Journal of Management & Social Sciences (ISSN 2455-2267), 11(3), 108. 10.21013/jmss.v11.n3.p1

Brand, C. M., & Marchant, L. F. (2015). Hair plucking in captive bonobos (*Pan paniscus*). Applied Animal Behaviour Science, 171, 192–196. 10.1016/j.applanim.2015.08.002

Brent, L., Koban, T., & Ramirez, S. (2002). Abnormal, abusive, and stress-related behaviors in baboon mothers. Biological Psychiatry, 52(11), 1047–1056. 10.1016/S0006-3223(02)01540-8

Bridges, R. S. (2015). Neuroendocrine regulation of maternal behavior. Frontiers in Neuroendocrinology, 36, 178–196. 10.1016/j.yfrne.2014.11.007

Burgener, N., Gusset, M., & Schmid, H. (2008). Frustrated appetitive foraging behavior, stereotypic pacing, and fecal glucocorticoid levels in snow leopards (*Uncia uncia*) in the Zurich zoo. Journal of Applied Animal Welfare Science, 11(1), 74–83. 10.1080/10888700701729254

Carini, L. M., Murgatroyd, C. A., & Nephew, B. C. (2013). Using chronic social stress to model postpartum depression in lactating rodents. Journal of Visualized Experiments : JoVE, 76, 50324. 10.3791/50324

Carlstead, K. (1998). Determining the Causes of Stereotypic Behaviors in Zoo Carnivores: Toward Appropriate Enrichment Strategies. 172–183.

Castles, D. L., Whiten, A., & Aureli, F. (1999). Social anxiety, relationships and self-directed behaviour among wild female olive baboons. Animal Behaviour, 58(6), 1207–1215. 10.1006/anbe.1999.1250

Cheney, D. L., & Seyfarth, R. M. (1999). Recognition of other individuals’ social relationships by female baboons. Animal Behaviour, 58(1), 67–75. 10.1006/anbe.1999.1131

Chu, B., Marwaha, K., Sanvictores, T., Awosika, A. O., & Ayers, D. (2025). Physiology, stress reaction. In StatPearls. StatPearls Publishing. http://www.ncbi.nlm.nih.gov/books/NBK541120/

Clark, S. 2021 Jun 4. Aye-aye. Duke Lemur Center. https://lemur.duke.edu/discover/meet-the-lemurs/aye-aye/

Clark, S. 2019. “Infant Announcement: Rare aye-aye born at the Duke Lemur Center”. Duke Lemur Center. https://lemur.duke.edu/melisandre/

Cless, I. T., Voss-Hoynes, H. A., Ritzmann, R. E., & Lukas, K. E. (2015). Defining pacing quantitatively: A comparison of gait characteristics between pacing and non-repetitive locomotion in zoo-housed polar bears. Applied Animal Behaviour Science, 169, 78–85. 10.1016/j.applanim.2015.04.002

Clinchy, M., Sheriff, M. J., & Zanette, L. Y. (2013). Predator-induced stress and the ecology of fear. Functional Ecology, 27(1), 56–65. 10.1111/1365-2435.12007

Coe, C. L., Kramer, M., Czéh, B., Gould, E., Reeves, A. J., Kirschbaum, C., & Fuchs, E. (2003). Prenatal stress diminishes neurogenesis in the dentate gyrus of juvenile rhesus monkeys. Biological Psychiatry, 54(10), 1025–1034. 10.1016/s0006-3223(03)00698-x

Coe, C. L., Lubach, G. R., & Shirtcliff, E. A. (2007). Maternal stress during pregnancy predisposes for iron deficiency in infant monkeys impacting innate immunity. Pediatric Research, *61*(5 Pt 1), 520–524. 10.1203/pdr.0b013e318045be53

Coe, C. L., & Shirtcliff, E. A. (2004). Growth trajectory evident at birth affects age of first delivery in female monkeys. Pediatric Research, 55(6), 914–920. 10.1203/01.PDR.0000125259.45025.4D

Coleman, K., & Pierre, P. J. (2014). Assessing anxiety in nonhuman primates. ILAR Journal, 55(2), 333–346. 10.1093/ilar/ilu019

Cooper, J. J., & Nicol, C. J. (1993). The “coping” hypothesis of stereotypic behaviour: A reply to Rushen. Animal Behaviour, 45(3), 616–618. 10.1006/anbe.1993.1072

Corder, K. M., Cortes, M. A., Bartley, A. F., Lear, S. A., Lubin, F. D., & Dobrunz, L. E. (2018). Prefrontal cortex-dependent innate behaviors are altered by selective knockdown of Gad1 in neuropeptide Y interneurons. PLOS ONE, 13(7), e0200809. 10.1371/journal.pone.0200809

Crockett, C. M., Sackett, G. P., Sandman, C. A., Chicz-DeMet, A., & Bentson, K. L. (2007). Beta-endorphin levels in longtailed and pigtailed macaques vary by abnormal behavior rating and sex. Peptides, 28(10), 1987–1997. 10.1016/j.peptides.2007.07.014

Dantzer, B., Newman, A. E. M., Boonstra, R., Palme, R., Boutin, S., Humphries, M. M., & McAdam, A. G. (2013). Density triggers maternal hormones that increase adaptive offspring growth in a wild mammal. *Science (New York*, N.Y.), 340(6137), 1215–1217. 10.1126/science.1235765

Daw, J. R., MacCallum-Bridges, C. L., & Admon, L. K. (2025). Trends and disparities in maternal self-reported mental and physical health. JAMA Internal Medicine, 185(7), 857. 10.1001/jamainternmed.2025.1260

Del Giudice, M., Buck, C. L., Chaby, L. E., Gormally, B. M., Taff, C. C., Thawley, C. J., Vitousek, M. N., & Wada, H. (2018). What is stress? A systems perspective. Integrative and Comparative Biology. 10.1093/icb/icy114

Demuru, E., Ferrari, P. F., & Palagi, E. (2018). Is birth attendance a uniquely human feature? New evidence suggests that Bonobo females protect and support the parturient. Evolution and Human Behavior, 39(5), 502–510. 10.1016/j.evolhumbehav.2018.05.003

Deputte, B. L. (1994). Ethological study of yawning in primates. I. Quantitative analysis and study of causation in two species of old world monkeys (*Cercocebus albigena* and *Macaca fascicularis*). Ethology, 98(3–4), 221–245. 10.1111/j.1439-0310.1994.tb01073.x

Deschamps, S., Woodside, B., & Walker, C. -D. (2003). Pups presence eliminates the stress hyporesponsiveness of early lactating females to a psychological stress representing a threat to the pups. Journal of Neuroendocrinology, 15(5), 486–497. 10.1046/j.1365-2826.2003.01022.x

Diezinger, F., & Anderson, J. R. (1986). Starting from scratch: A first look at a “displacement activity” in group-living rhesus monkeys. American Journal of Primatology, 11(2), 117–124. 10.1002/ajp.1350110204

Downey, C., & Crummy, A. (2022). The impact of childhood trauma on children’s wellbeing and adult behavior. European Journal of Trauma & Dissociation, 6(1), 100237. 10.1016/j.ejtd.2021.100237

Duke Lemur Center. (2016). Aye-aye husbandry guidelines. https://lemur.duke.edu/accordions/aye-aye-husbandry-guidelines/

Dunayer, E. S., & Berman, C. M. (2018). Infant handling among primates. International Journal of Comparative Psychology, 31(0). 10.46867/ijcp.2018.31.02.06

Easley, S. P., Coelho, A. M., & Taylor, L. L. (1987). Scratching, dominance, tension, and displacement in male baboons. American Journal of Primatology, 13(4), 397–411. 10.1002/ajp.1350130405

Elias, M. J. (1989). Schools as a source of stress to children: An analysis of causal and ameliorative influences. Journal of School Psychology, 27(4), 393–407. 10.1016/0022-4405(89)90016-2

Erickson, C. J. (1991). Percussive foraging in the aye-aye, *Daubentonia madagascariensis*. Animal Behaviour, 41(5), 793–801. 10.1016/S0003-3472(05)80346-X

Erickson, C. J. (1995). Feeding sites for extractive foraging by the aye-aye, *Daubentonia madagascariensis*. American Journal of Primatology, 35(3), 235–240. 10.1002/ajp.1350350306

Feistner, A. T. C., & Ashbourne, C. J. (1994). Infant development in a captive-bred aye-aye (*Daubentonia madagascariensis*) over the first year of life. Folia Primatologica, 62(1–3), 74–92. 10.1159/000156765

Feldman, R., Singer, M., & Zagoory, O. (2010). Touch attenuates infants’ physiological reactivity to stress. Developmental Science, 13(2), 271–278. 10.1111/j.1467-7687.2009.00890.x

Fernandez, E. J. (2021). Appetitive search behaviors and stereotypies in polar bears (*Ursus maritimus*). Behavioural Processes, 182, 104299. 10.1016/j.beproc.2020.104299

Ferrante, A., & Ferrante, A. (2015). Il problema del succhiamento del dito. Nuove interpretazioni e implicazioni terapeutiche [Finger or thumb sucking. New interpretations and therapeutic implications]. Minerva Pediatrica, 67(4), 285–297. PMID: 26129804.

Feusner, J., Hembacher, E., & Phillips, K. A. (2009). The mouse who couldn’t stop washing: Pathologic grooming in animals and humans. CNS Spectrums, 14(9), 503–513. 10.1017/s1092852900023567

Finlay-Jones, R., & Brown, G. W. (1981). Types of stressful life event and the onset of anxiety and depressive disorders. Psychological Medicine, 11(4), 803–815. 10.1017/S0033291700041301

Fisher, R. A. (1935). The logic of inductive inference. Journal of the Royal Statistical Society, 98(1), 39–54. 10.2307/2342435

Fisher, R. A. (2018). On the interpretation of χ2 from contingency tables, and the calculation of p. Journal of the Royal Statistical Society Series A (Statistics in Society), 85(1), 87–94. 10.1111/j.2397-2335.1922.tb00768.x

Fleming, A. S., & Anderson, V. (1987). Affect and nurturance: Mechanisms mediating maternal behavior in two female mammals. Progress in Neuro-Psychopharmacology & Biological Psychiatry, 11(2–3), 121–127. 10.1016/0278-5846(87)90049-2

Fleming, A. S., Steiner, M., & Corter, C. (1997). Cortisol, hedonics, and maternal responsiveness in human mothers. Hormones and Behavior, 32(2), 85–98. 10.1006/hbeh.1997.1407

Fox, M. W. (1984). Farm Animals: Husbandry, Behavior and Veterinary Practice. (p. 184). Baltimore: University Park Press.

Friedman, M. (1937). The use of ranks to avoid the assumption of normality implicit in the analysis of variance. Journal of the American Statistical Association, 32(200), 675–701. 10.1080/01621459.1937.10503522

Fruth, B., & Hohmann, G. (1996). Nest building behavior in the great apes: The great leap forward? In W. C. McGrew, L. F. Marchant, & T. Nishida (Eds.), Great Ape Societies (1st ed., pp. 225–240). Cambridge University Press. 10.1017/CBO9780511752414.019

Gallup, A. C. (2022). The causes and consequences of yawning in animal groups. Animal Behaviour, 187, 209–219. 10.1016/j.anbehav.2022.03.011

Gaynor, K. M., & Cords, M. (2012). Antipredator and social monitoring functions of vigilance behaviour in blue monkeys. Animal Behaviour, 84(3), 531–537. 10.1016/j.anbehav.2012.06.003

Global Burden of Disease (GBD). (2021). Seattle, United States: Institute for Health Metrics and Evaluation (IHME); 2024 (https://vizhub.healthdata.org/gbd-results/)

Gluckman, P. D., & Hanson, M. A. (2004). Living with the past: Evolution, development, and patterns of disease. Science, 305(5691), 1733–1736. 10.1126/science.1095292

Goldberg, M. B., Langman, V. A., & Richard Taylor, C. (1981). Panting in dogs: Paths of air flow in response to heat and exercise. Respiration Physiology, 43(3), 327–338. 10.1016/0034-5687(81)90113-4

Goodall, J., & Athumani, J. (1980). An observed birth in a free-living chimpanzee (*Pan troglodytes schweinfurthii*) in Gombe National Park, Tanzania. Primates, 21(4), 545–549. 10.1007/BF02373843

Grigg, E. K., Chou, J., Parker, E., Gatesy-Davis, A., Clarkson, S. T., & Hart, L. A. (2021). Stress-related behaviors in companion dogs exposed to common household noises, and owners’ interpretations of their dogs’ behaviors. Frontiers in Veterinary Science, 8, 760845. 10.3389/fvets.2021.760845

Gupta, S., & Mittal, S. (2013). Yawning and its physiological significance. International Journal of Applied and Basic Medical Research, 3(1), 11. 10.4103/2229-516X.112230

Gust, D. A., Gordon, T. P., Hambright, M. K., & Wilson, M. E. (1993). Relationship between social factors and pituitary-adrenocortical activity in female rhesus monkeys (*Macaca mulatta*). Hormones and Behavior, 27(3), 318–331. 10.1006/hbeh.1993.1024

Hansen, E. W. (1966). The Development of Maternal and Infant Behavior in the Rhesus Monkey. Behaviour, 27(1/2), 107–149. http://www.jstor.org/stable/4533153

Harb, F., Liuzzi, M. T., Huggins, A. A., Webb, E. K., Fitzgerald, J. M., Krukowski, J. L., deRoon-Cassini, T. A., & Larson, C. L. (2024). Childhood maltreatment and amygdala-mediated anxiety and posttraumatic stress following adult trauma. Biological Psychiatry Global Open Science, 4(4), 100312. 10.1016/j.bpsgos.2024.100312

Hartstone-Rose, A., Dickinson, E., Boettcher, M. L., & Herrel, A. (2020). A primate with a Panda’s thumb: The anatomy of the pseudothumb of *Daubentonia madagascariensis*. American Journal of Physical Anthropology, 171(1), 8–16. 10.1002/ajpa.23936

Heagerty, A., Wales, R. A., Prongay, K., Gottlieb, D. H., & Coleman, K. (2017). Social hair pulling in captive rhesus macaques (*Macaca mulatta*). American Journal of Primatology, 79(12), 10.1002/ajp.22720. 10.1002/ajp.22720

Hock, E., McBride, S., & Gnezda, M. T. (1989). Maternal separation anxiety: Mother-infant separation from the maternal perspective. Child Development, 60(4), 793. 10.2307/1131019

Hoffman, C. L., Ayala, J. E., Mas-Rivera, A., & Maestripieri, D. (2010). Effects of reproductive condition and dominance rank on cortisol responsiveness to stress in free-ranging female rhesus macaques. American Journal of Primatology, 72(7), 559–565. 10.1002/ajp.20793

Holley, J. M., & Simpson, M. J. A. (1981). A comparison of primiparous and multiparous mother-infant dyads in *Macaca mulatta*. Primates, 22(3), 379–392. 10.1007/BF02381578

Hühne, A., Volkmann, P., Stephan, M., Rossner, M., & Landgraf, D. (2020). An in-depth neurobehavioral characterization shows anxiety-like traits, impaired habituation behavior, and restlessness in male *Cryptochrome* -deficient mice. *Genes*, Brain and Behavior, 19(8), e12661. 10.1111/gbb.12661

IUCN. 2025. The IUCN Red List of Threatened Species. Version 2025-2. https://www.iucnredlist.org.

Jeličić, L., Veselinović, A., Ćirović, M., Jakovljević, V., Raičević, S., & Subotić, M. (2022). Maternal distress during pregnancy and the postpartum period: Underlying mechanisms and child’s developmental outcomes—a narrative review. International Journal of Molecular Sciences, 23(22), 13932. 10.3390/ijms232213932

Kano, T. (1979). A pilot study on the ecology of pygmy chimpanzees, Pan paniscus. The Great Apes, 5, 123–135.

Karayağız, Ş., Aktan, T., & Karayağız, L. Z. (2020). Parental attachment patterns in mothers of children with anxiety disorder. Children, 7(5), 46. 10.3390/children7050046

Kaufman, J. A., Ahrens, E. T., Laidlaw, D. H., Zhang, S., & Allman, J. M. (2005). Anatomical analysis of an aye-aye brain (*Daubentonia madagascariensis*, primates: Prosimii) combining histology, structural magnetic resonance imaging, and diffusion-tensor imaging. *The Anatomical Record Part A: Discoveries in Molecular*, Cellular, and Evolutionary Biology, 287A(1), 1026–1037. 10.1002/ar.a.20264

Kay, R. F., & Kirk, E. C. (2000). Osteological evidence for the evolution of activity pattern and visual acuity in primates. American Journal of Physical Anthropology, 113(2), 235–262. 10.1002/1096-8644(200010)113:2%3C235::AID-AJPA7%3E3.0.CO;2-9

Kemps, A., & Timmermans, P. (1982). Parturition behaviour in pluriparous Java-macaques (*Macaca fascicularis*). Primates, 23(1), 75–88. 10.1007/BF02381439

Kendler, K. S., Karkowski, L. M., & Prescott, C. A. (1999). Causal relationship between stressful life events and the onset of major depression. American Journal of Psychiatry, 156(6), 837–841. 10.1176/ajp.156.6.837

Kennes, D., Ödberg, F. O., Bouquet, Y., & De Rycke, P. H. (1988). Changes in naloxone and haloperidol effects during the development of captivity-induced jumping stereotypy in bank voles. European Journal of Pharmacology, 153(1), 19–24. 10.1016/0014-2999(88)90583-3

Keverne, E. B., Martensz, N. D., & Tuite, B. (1989). Beta-endorphin concentrations in cerebrospinal fluid of monkeys are influenced by grooming relationships. Psychoneuroendocrinology, 14(1–2), 155–161. 10.1016/0306-4530(89)90065-6

Kinnally, E. L., Martinez, S. J., Chun, K., Capitanio, J. P., & Ceniceros, L. C. (2019). Early social stress promotes inflammation and disease risk in rhesus monkeys. Scientific Reports, 9(1), 7609. 10.1038/s41598-019-43750-1

Klopfer, P. H., & Dugard, J. (1976). Patterns of maternal care in lemurs: III. Lemur variegatus. Zeitschrift Für Tierpsychologie, 40(2), 210–220. 10.1111/j.1439-0310.1976.tb00933.x

Knezevic, E., Nenic, K., Milanovic, V., & Knezevic, N. N. (2023). The role of cortisol in chronic stress, neurodegenerative diseases, and psychological disorders. Cells, 12(23), 2726. 10.3390/cells12232726

Krpan, K. M., Coombs, R., Zinga, D., Steiner, M., & Fleming, A. S. (2005). Experiential and hormonal correlates of maternal behavior in teen and adult mothers. Hormones and Behavior, 47(1), 112–122. 10.1016/j.yhbeh.2004.08.006

Kutsukake, N. (2006). The context and quality of social relationships affect vigilance behaviour in wild chimpanzees. Ethology, 112(6), 581–591. 10.1111/j.1439-0310.2006.01200.x

Laméris, D. W., Verspeek, J., Salas, M., Staes, N., Torfs, J. R. R., Eens, M., & Stevens, J. M. G. (2022). Evaluating self-directed behaviours and their association with emotional arousal across two cognitive tasks in bonobos(Pan paniscus). Animals : An Open Access Journal from MDPI, 12(21), 3002. 10.3390/ani12213002

Lee, T. M., & Zucker, I. (1988). Vole infant development is influenced perinatally by maternal photoperiodic history. *American Journal of Physiology-Regulatory*, Integrative and Comparative Physiology, 255(5), R831–R838. 10.1152/ajpregu.1988.255.5.R831

Leone, A., Ferrari, P. F., & Palagi, E. (2014). Different yawns, different functions? Testing social hypotheses on spontaneous yawning in *Theropithecus gelada*. Scientific Reports, 4(1), 4010. 10.1038/srep04010

Lévy, F. (2016). Neuroendocrine control of maternal behavior in non-human and human mammals. Annales d’Endocrinologie, 77(2), 114–125. 10.1016/j.ando.2016.04.002

Liu, S., Tian, H., Ren, S., Sun, W., Fan, P., Xia, D., Sun, B., Li, J., & Wang, X. (2024). Social risk to infant: The role of kin for maternal visual monitoring in Tibetan macaques. Ecology and Evolution, 14(6), e11626. 10.1002/ece3.11626

Lopez-Vergara, L., Santillan-Doherty, A. M., Mayagoitia, L., & Mondragon-Ceballos, R. (1989). Self and social grooming in stump-tailed macaques: Effects of kin presence or absence within the group. Behavioural Processes, 18(1), 99–106. 10.1016/S0376-6357(89)80008-7

Louis Jr, E.E., Sefczek, T.M., Randimbiharinirina, D.R., Raharivololona, B., Rakotondrazandry, J.N., Manjary, D., Aylward, M. & Ravelomandrato, F. (2020). Daubentonia madagascariensis. The IUCN Red List of Threatened Species 2020: e.T6302A115560793. 10.2305/IUCN.UK.2020-2.RLTS.T6302A115560793.en.

Love, O. P., McGowan, P. O., & Sheriff, M. J. (2013). Maternal adversity and ecological stressors in natural populations: The role of stress axis programming in individuals, with implications for populations and communities. Functional Ecology, 27(1), 81–92. 10.1111/j.1365-2435.2012.02040.x

Lutz, C., Well, A., & Novak, M. (2003). Stereotypic and self-injurious behavior in rhesus macaques: A survey and retrospective analysis of environment and early experience. American Journal of Primatology, 60(1), 1–15. 10.1002/ajp.10075

MacIntosh, A. J. J., & Sicotte, P. (2009). Vigilance in ursine black and white colobus monkeys (*Colobus vellerosus*): An examination of the effects of conspecific threat and predation. American Journal of Primatology, 71(11), 919–927. 10.1002/ajp.20730

Maestripieri, D. (1993a). Maternal anxiety in rhesus macaques (*Macaca mulatta*): I. Measurement of anxiety and identification of anxiety-eliciting situations. Ethology, 95(1), 19–31. 10.1111/j.1439-0310.1993.tb00453.x

Maestripieri, D. (1993b). Vigilance costs of allogrooming in macaque mothers. The American Naturalist, 141(5), 744–753. 10.1086/285503

Maestripieri, D. (1995). First steps in the macaque world: Do rhesus mothers encourage their infants’ independent locomotion? Animal Behaviour, 49(6), 1541–1549. 10.1016/0003-3472(95)90075-6

Maestripieri, D. (1996). Maternal encouragement of infant locomotion in pigtail macaques, *Macaca nemestrina*. Animal Behaviour, 51(3), 603–610. 10.1006/anbe.1996.0064

Maestripieri, D. (2001). Intraspecific variability in parenting styles of rhesus macaques (*Macaca mulatta*): The role of the social environment. Ethology, 107(3), 237–248. 10.1046/j.1439-0310.2001.00661.x

Maestripieri, D. (2005). Effects of early experience on female behavioural and reproductive development in rhesus macaques. Proceedings of the Royal Society B: Biological Sciences, 272(1569), 1243–1248. 10.1098/rspb.2005.3059

Maestripieri, D. (2011). Emotions, stress, and maternal motivation in primates. American Journal of Primatology, 73(6), 516–529. 10.1002/ajp.20882

Maestripieri, D., Schino, G., Aureli, F., & Troisi, A. (1992). A modest proposal: Displacement activities as an indicator of emotions in primates. Animal Behaviour, 44(5), 967–979. 10.1016/S0003-3472(05)80592-5

Maestripieri, D., Hoffman, C. L., Anderson, G. M., Carter, C. S., & Higley, J. D. (2009). Mother–infant interactions in free-ranging rhesus macaques: Relationships between physiological and behavioral variables. Physiology & Behavior, 96(4–5), 613–619. 10.1016/j.physbeh.2008.12.016

Marais, L., Daniels, W., Brand, L., Viljoen, F., Hugo, C., & Stein, D. J. (2006). Psychopharmacology of maternal separation anxiety in vervet monkeys. Metabolic Brain Disease, 21(2–3), 191–200. 10.1007/s11011-006-9011-8

Marks, I. M., & Nesse, R. M. (1994). Fear and fitness: An evolutionary analysis of anxiety disorders. Ethology and Sociobiology, 15(5), 247–261. 10.1016/0162-3095(94)90002-7

Maródi, L. (2006). Neonatal innate immunity to infectious agents. Infection and Immunity, 74(4), 1999–2006. 10.1128/IAI.74.4.1999-2006.2006

Marriner, L. M., & Drickamer, L. C. (1994). Factors influencing stereotyped behavior of primates in a zoo. Zoo Biology, 13(3), 267–275. 10.1002/zoo.1430130308

Martin, P. R., & Bateson, P. (2007). Measuring behaviour: An introductory guide (3rd ed). Cambridge University Press.

Mason, G. J. (1991). Stereotypies: A critical review. Animal Behaviour, 41(6), 1015–1037. 10.1016/S0003-3472(05)80640-2

Mathew, J., & Paulose, C. S. (2011). The healing power of well-being. Acta Neuropsychiatrica, 23(4), 145–155. 10.1111/j.1601-5215.2011.00578.x

McCormack, K., Howell, B. R., Guzman, D., Villongco, C., Pears, K., Kim, H., Gunnar, M. R., & Sanchez, M. M. (2015). The development of an instrument to measure global dimensions of maternal care in rhesus macaques (*Macaca mulatta*). American Journal of Primatology, 77(1), 20–33. 10.1002/ajp.22307

McDougall, P. (2011). Scratching our heads: Rethinking social anxiety in vervets (*Chlorocebus aethiops*). International Journal of Primatology, 32(2), 335–345. 10.1007/s10764-010-9471-x

McEwen, B. S. (1998). Protective and damaging effects of stress mediators. New England Journal of Medicine, 338(3), 171–179. 10.1056/NEJM199801153380307

McKee, K., Admon, L. K., Winkelman, T. N. A., Muzik, M., Hall, S., Dalton, V. K., & Zivin, K. (2020). Perinatal mood and anxiety disorders, serious mental illness, and delivery-related health outcomes, United States, 2006–2015. BMC Women’s Health, 20(1), 150. 10.1186/s12905-020-00996-6

Mishra, A. K., & Varma, A. R. (2023). A comprehensive review of the generalized anxiety disorder. Cureus. 10.7759/cureus.46115

Mitchell, G., & Gomber, J. (1976). Moving laboratory rhesus monkeys (*Macaca mulatta*) to unfamiliar home cages. Primates, 17(4), 543–546. 10.1007/BF02382913

Moberg, G. P., & Mench, J. A. (Eds.). (2000). *The biology of animal stress: Basic principles and implications for animal welfare*. (1st ed.). (p. 3). CABI Publishing. 10.1079/9780851993591.0000

Mota-Rojas, D., Orihuela, A., Strappini, A., Villanueva-García, D., Napolitano, F., Mora-Medina, P., Barrios-García, H. B., Herrera, Y., Lavalle, E., & Martínez-Burnes, J. (2020). Consumption of maternal placenta in humans and nonhuman mammals: Beneficial and adverse effects. Animals, 10(12), 2398. 10.3390/ani10122398

Nguyen, N., Gesquiere, L. R., Wango, E. O., Alberts, S. C., & Altmann, J. (2008). Late pregnancy glucocorticoid levels predict responsiveness in wild baboon mothers (*Papio cynocephalus*). Animal Behaviour, 75(5), 1747–1756. 10.1016/j.anbehav.2007.09.035

Nguyen, N., Lee, L. M., Fashing, P. J., Nurmi, N. O., Stewart, K. M., Turner, T. J., Barry, T. S., Callingham, K. R., Goodale, C. B., Kellogg, B. S., Burke, R. J., Bechtold, E. K., Claase, M. J., Eriksen, G. A., Jones, S. C. Z., Kerby, J. T., Kraus, J. B., Miller, C. M., Trew, T. H., … Venkataraman, V. V. (2017). Comparative primate obstetrics: Observations of 15 diurnal births in wild gelada monkeys (*Theropithecus gelad*a) and their implications for understanding human and nonhuman primate birth evolution. American Journal of Physical Anthropology, 163(1), 14–29. 10.1002/ajpa.23141

Ninan, P. T., Insel, T. M., Cohen, R. M., Cook, J. M., Skolnick, P., & Paul, S. M. (1982). Benzodiazepine receptor-mediated experimental “anxiety” in primates. Science, 218(4579), 1332–1334. 10.1126/science.6293059

Norscia, I., Agostini, L., Moroni, A., Caselli, M., Micheletti-Cremasco, M., Vardé, C., & Palagi, E. (2021). Yawning is more contagious in pregnant than nulliparous women: Naturalistic and experimental evidence. Human Nature, 32(2), 301–325. 10.1007/s12110-021-09404-w

Nowak, R. (2000). Role of mother-young interactions in the survival of offspring in domestic mammals. Reviews of Reproduction, 5(3), 153–163. 10.1530/ror.0.0050153.

Ocepek, M., & Andersen, I. L. (2018). Sow communication with piglets while being active is a good predictor of maternal skills, piglet survival and litter quality in three different breeds of domestic pigs (*Sus scrofa domesticus*). PLOS ONE, 13(11), e0206128. 10.1371/journal.pone.0206128

Overall, K. L., Dunham, A. E., & Frank, D. (2001). Frequency of nonspecific clinical signs in dogs with separation anxiety, thunderstorm phobia, and noise phobia, alone or in combination. Journal of the American Veterinary Medical Association, 219(4), 467–473. 10.2460/javma.2001.219.467

Owen, R. (1866). On the aye-aye (*Chiromys madagascarensis*). Trans Zool Soc Lond, 5, 33–101.

Palagi, E., Leone, A., Mancini, G., & Ferrari, P. F. (2009). Contagious yawning in gelada baboons as a possible expression of empathy. Proceedings of the National Academy of Sciences, 106(46), 19262–19267. 10.1073/pnas.0910891106

Palagi, E., & Norscia, I. (2011). Scratching around stress: Hierarchy and reconciliation make the difference in wild brown lemurs (*Eulemur fulvus*). Stress, 14(1), 93–97. 10.3109/10253890.2010.505272

Paukner, A., & Anderson, J. R. (2006). Video-induced yawning in stumptail macaques (*Macaca arctoides*). Biology Letters, 2(1), 36–38. 10.1098/rsbl.2005.0411

Petter, J. & Peyrieras, A. (1970). Nouvelle contribution a l’etude d’un lemurien Malgache, le aye-aye [New contribution to the study of a Malagasy lemuran, the aye-aye]. (Daubentonia madagascariensis E. Geoggrey). Mammalia, 34(2), 167–193. 10.1515/mamm.1970.34.2.167

Philbin, N. (1998). Towards an understanding of stereotypic behaviour in laboratory macaques. Animal Welfare Institute.

https://awionline.org/content/towards-understanding-stereotypic-behaviour-laboratory-macaques

Poirier, C., Oliver, C. J., Castellano Bueno, J., Flecknell, P., & Bateson, M. (2019). Pacing behaviour in laboratory macaques is an unreliable indicator of acute stress. Scientific Reports, 9(1), 7476. 10.1038/s41598-019-43695-5

Quinn, A., & Wilson, D. E. (2004). Daubentonia madagascariensis. Mammalian Species, 740, 1–6. 10.1644/740

Radoš, S. N., Tadinac, M., & Herman, R. (2018). Anxiety during pregnancy and postpartum: Course, predictors and comorbidity with postpartum depression. Acta Clinica Croatica, 57(1), 39–51. 10.20471/acc.2018.57.01.05

Rakotondrazandry, J. N., Ravelomandrato, F., Sefczek, T. M., Andriamalala, Y. R., Frasier, C. L., Villanova, V. L., Rasoloharijaona, S., Raveloson, H., & Louis, E. E. (2021). Developmental timeline of wild aye-aye (*Daubentonia madagascariensis*) infants in Kianjavato and Torotorofotsy, Madagascar. International Journal of Primatology, 42(3), 344–348. 10.1007/s10764-021-00216-4

Reamer, L., Tooze, Z., Coulson, C., & Semple, S. (2010). Correlates of self-directed and stereotypic behaviours in captive red-capped mangabeys (*Cercocebus torquatus torquatus*). Applied Animal Behaviour Science, 124(1–2), 68–74. 10.1016/j.applanim.2010.01.012

Redmond, D. E., & Huang, Y. H. (1979). II. New evidence for a locus coeruleus-norepinephrine connection with anxiety. Life Sciences, 25(26), 2149–2162. 10.1016/0024-3205(79)90087-0

Reinhardt, V. (2005). Hair pulling: A review. Laboratory Animals, 39(4), 361–369. 10.1258/002367705774286448

Rushen, J. (1993). The “coping” hypothesis of stereotypic behaviour. Animal Behaviour, 45(3), 613–615. 10.1006/anbe.1993.1071

Ryu, S., & Fan, L. (2023). The relationship between financial worries and psychological distress among U.S. Adults. Journal of Family and Economic Issues, 44(1), 16–33. 10.1007/s10834-022-09820-9

Salali, G. D., Uysal, M. S., & Bevan, A. (2021). Adaptive function and correlates of anxiety during a pandemic. Evolution, Medicine, and Public Health, 9(1), 393–405. 10.1093/emph/eoab037

Saltzman, W., & Abbott, D. H. (2009). Effects of elevated circulating cortisol concentrations on maternal behavior in common marmoset monkeys (*Callithrix jacchus*). Psychoneuroendocrinology, 34(8), 1222–1234. 10.1016/j.psyneuen.2009.03.012

Sanchez, M. M., Mccormack, K., Grand, A. P., Fulks, R., Graff, A., & Maestripieri, D. (2010). Effects of sex and early maternal abuse on adrenocorticotropin hormone and cortisol responses to the corticotropin-releasing hormone challenge during the first 3 years of life in group-living rhesus monkeys. Development and Psychopathology, 22(1), 45–53. 10.1017/S0954579409990253

Schino, G., Perretta, G., Taglioni, A. M., Monaco, V., & Troisi, A. (1996). Primate displacement activities as an ethopharmacological model of anxiety. Anxiety, 2(4), 186–191. 10.1002/(SICI)1522-7154(1996)2:4<186::AID-ANXI5>3.0.CO;2-M

Schino, G., Speranza, L., Ventura, R., & Troisi, A. (2003). Infant handling and maternal response in Japanese macaques. International Journal of Primatology, 24(3), 627–638. 10.1023/A:1023796531972

Schino, G., Troisi, A., Perretta, G., & Monaco, V. (1991). Measuring anxiety in nonhuman primates: Effect of lorazepam on macaque scratching. Pharmacology Biochemistry and Behavior, 38(4), 889–891. 10.1016/0091-3057(91)90258-4

Schmidt-Nielsen, K., Hainsworth, F. R., & Murrish, D. E. (1970). Counter-current heat exchange in the respiratory passages: Effect on water and heat balance. Respiration Physiology, 9(2), 263–276. 10.1016/0034-5687(70)90075-7

Sclafani, V., Norscia, I., Antonacci, D., & Palagi, E. (2012). Scratching around mating: Factors affecting anxiety in wild Lemur catta. Primates, 53(3), 247–254. 10.1007/s10329-012-0294-6

Seraphin, S. B., Sanchez, M. M., Whitten, P. L., & Winslow, J. T. (2022). The behavioral neuroendocrinology of dopamine systems in differently reared juvenile male rhesus monkeys (*Macaca mulatta*). Hormones and Behavior, 137, 105078. 10.1016/j.yhbeh.2021.105078

Short, J. L. (2002). The effects of parental divorce during childhood on college students. Journal of Divorce & Remarriage, 38(1–2), 143–155. 10.1300/J087v38n01_08

Silk, J. B., Rendall, D., Cheney, D. L., & Seyfarth, R. M. (2003). Natal attraction in adult female baboons (*Papio cynocephalus ursinus*) in the Moremi Reserve, Botswana. Ethology, 109(8), 627–644. 10.1046/j.1439-0310.2003.00907.x

Simmons, D., & Self, D. W. (2009). Role of mu- and delta-opioid receptors in the nucleus accumbens in cocaine-seeking behavior. Neuropsychopharmacology, 34(8), 1946–1957. 10.1038/npp.2009.28

Solanki, G.S., & Zothansiama. (2013). Births in Captive Stump-Tailed Macaques (*Macaca arctoides*). Folia Primatologica, 84(6), 394–404. 10.1159/000353171

Soligo, C. (2005). Anatomy of the hand and arm in *Daubentonia madagascariensis*: A functional and phylogenetic outlook. Folia Primatologica, 76(5), 262–300. 10.1159/000088034

Spruijt, B. M., van Hooff, J. A., & Gispen, W. H. (1992). Ethology and neurobiology of grooming behavior. Physiological Reviews, 72(3), 825–852. 10.1152/physrev.1992.72.3.825

Stanton, M. A., Heintz, M. R., Lonsdorf, E. V., Santymire, R. M., Lipende, I., & Murray, C. M. (2015). Maternal behavior and physiological stress levels in wild chimpanzees (*Pan troglodytes schweinfurthii*). International Journal of Primatology, 36(3), 473–488. 10.1007/s10764-015-9836-2

Steenbeek, R., Piek, R. C., van Buul, M., & van Hooff, J. A. R. A. M. (1999). Vigilance in wild Thomas’s langurs (*Presbytis thomasi*): The importance of infanticide risk. Behavioral Ecology and Sociobiology, 45(2), 137–150. 10.1007/s002650050547

Steimer, T. (2002). The biology of fear- and anxiety-related behaviors. Dialogues in Clinical Neuroscience, 4(3), 231–249. 10.31887/DCNS.2002.4.3/tsteimer

Stewart, F. A., & Pruetz, J. D. (2013). Do chimpanzee nests serve an anti-predatory function? American Journal of Primatology, 75(6), 593–604. 10.1002/ajp.22138

Stewart, F. A., Pruetz, J. D., & Hansell, M. H. (2007). Do chimpanzees build comfortable nests? American Journal of Primatology, 69(8), 930–939. 10.1002/ajp.20432

Struhsaker, T. T. (1967). Social Structure among Vervet Monkeys (*Cercopithecus aethiops*). Behaviour, 29(2/4), 83–121. http://www.jstor.org/stable/4533186

Tanaka, I., & Takefushi, H. (1993). Elimination of external parasites (Lice) is the primary function of grooming in free-ranging Japanese macaques. Anthropological Science, 101(2), 187–193. 10.1537/ase.101.187

Tatemoto, P., Broom, D. M., & Zanella, A. J. (2022). Changes in stereotypies: Effects over time and over generations. Animals, 12(19), 2504. 10.3390/ani12192504

Tecot, S. R., Baden, A. L., Romine, N. K., & Kamilar, J. M. (2012). Infant parking and nesting, not allomaternal care, influence Malagasy primate life histories. Behavioral Ecology and Sociobiology, 66(10), 1375–1386. 10.1007/s00265-012-1393-5

Tiira, K., Sulkama, S., & Lohi, H. (2016). Prevalence, comorbidity, and behavioral variation in canine anxiety. Journal of Veterinary Behavior, 16, 36–44. 10.1016/j.jveb.2016.06.008

Tinklepaugh, O. L., & Hartman, C. G. (1930). Behavioral aspects of parturition in the monkey (*Macacus rhesus*). Journal of Comparative Psychology, 11(1), 63–98. 10.1037/h0070309

Thompson, C. L., & Hermann, E. A. (2024). Behavioral thermoregulation in primates: A review of literature and future avenues. American Journal of Primatology, 86(6), e23614. 10.1002/ajp.23614

Treves, A., Drescher, A., & Snowdon, C. T. (2003). Maternal watchfulness in black howler monkeys (*Alouatta pigra*). Ethology, 109(2), 135–146. 10.1046/j.1439-0310.2003.00853.x

Troisi, A. (2002). Displacement activities as a behavioral measure of stress in nonhuman primates and human subjects. Stress, 5(1), 47–54. 10.1080/102538902900012378

Troisi, A., & Schino, G. (1987). Environmental and social influences on autogrooming behaviour in a captive group of Java monkeys. Behaviour. 10.1163/156853987X00161

Troisi, A., Schino, G., D’Antoni, M., Pandolfi, N., Aureli, F., & D’Amato, F. R. (1991). Scratching as a behavioral index of anxiety in macaque mothers. Behavioral and Neural Biology, 56(3), 307–313. 10.1016/0163-1047(91)90469-7

Turner, S. E., Fedigan, L. M., Nakamichi, M., Matthews, H. D., McKenna, K., Nobuhara, H., Nobuhara, T., & Shimizu, K. (2010). Birth in Free-ranging *Macaca fuscata*. International Journal of Primatology, 31(1), 15–37. 10.1007/s10764-009-9376-8

Van Lawick-Goodall, J. (1968). The behaviour of free-living chimpanzees in the Gombe stream reserve. Animal Behaviour Monographs, 1, 161-IN12. 10.1016/S0066-1856(68)80003-2

Ventura, R., Majolo, B., Schino, G., & Hardie, S. (2005). Differential effects of ambient temperature and humidity on allogrooming, self-grooming, and scratching in wild Japanese macaques. American Journal of Physical Anthropology, 126(4), 453–457. 10.1002/ajpa.20125

Vick, S., & Paukner, A. (2010). Variation and context of yawns in captive chimpanzees (*Pan troglodytes*). American Journal of Primatology, 72(3), 262–269. 10.1002/ajp.20781

Wang, M., & Saudino, K. J. (2011). Emotion regulation and stress. Journal of Adult Development, 18(2), 95–103. 10.1007/s10804-010-9114-7

Willott, J. F., & McDaniel, J. (1974). Changes in the behavior of laboratory-reared rhesus monkeys following the threat of separation. Primates, 15(4), 321–326. 10.1007/BF01791669

Woods, D. W., & Houghton, D. C. (2014). Diagnosis, evaluation, and management of trichotillomania. Psychiatric Clinics of North America, 37(3), 301–317. 10.1016/j.psc.2014.05.005

Zar, J. H. (1999). *Biostatistical analysis* (4. ed., internat. ed). Prentice Hall International. ZIMS. 2019. Species360 Zoological Information Management System (ZIMS). http://zims.Species360.org.

Zhang, Z., Li, N., Chen, R., Lee, T., Gao, Y., Yuan, Z., Nie, Y., & Sun, T. (2021). Prenatal stress leads to deficits in brain development, mood related behaviors and gut microbiota in offspring. Neurobiology of Stress, 15, 100333. 10.1016/j.ynstr.2021.100333

Zhao, H., Wang, Y., Li, J., Li, N., Zhou, W., Wang, C., & Li, B. (2025). Golden snub-nosed monkeys: Potential primate paradigm in post-traumatic stress disorder research. Biology, 14(2), 156. 10.3390/biology14020156

