## Supplementary Material for "First study of an aye-aye (*Daubentonia madagascariensis*) mother’s anxiety behaviors peripartum"

### Supplemental Figures

**Table S1**: Chi-Square test of the anxiety-related behavioral frequency of a mother aye-aye (n=1) peripartum (Day before birth: June 6; Day of birth: June 7; Day after birth: June 8). Significance value adjusted for Bonferroni correction ($\alpha = 0.0045$).

| Day | $\chi^2$ | df | p-value |
|---|---|---|---|
| June 6 | 1100.05 | 10 | < 0.001 |
| June 8 | 496.01 | 10 | < 0.001 |
| June 8 | 162.51 | 10 | < 0.001 |

**Table S2.** Chi-square goodness of fit examining the frequency of one anxiety-related behavior across three days peripartum (Day before birth: June 6; Day of birth: June 7; Day after birth: June 8) in a mother aye-aye (n=1). Significance value adjusted for Bonferroni correction **(**$\alpha$ = 0.0167).

| Behavior | $\chi^2$ | df | p-value |
|---|---|---|---|
| Auto (self) grooming | 30.34 | 2 | $p < 0.001$ |
| Auto (self) scratching | 112.93 | 2 | $p < 0.001$ |
| Examining Genitalia | 62.59 | 2 | $p < 0.001$ |
| Hyper-vigilance | 23.53 | 2 | $p < 0.001$ |
| Nest Construction | 337.4 | 2 | $p < 0.001$ |
| Nudging infant | 28 | 2 | $p < 0.001$ |
| Pacing | 5.43 | 2 | $p = 0.066$ |
| Shaking | 12 | 2 | $p < 0.001$ |
| Yawning | 6 | 2 | $p = 0.050$ |

**Table S3.** Pairwise day-by-day Chi square tests compare the frequency of behaviors across observation days (Day before birth: June 6; Day of birth: June 7; Day after birth: June 8) in a mother aye-aye (n=1). Significance was evaluated using a Bonferroni-corrected $\alpha = 0.0167$.

| Behavior | Comparison | $\chi^2$ | df | p-value |
|---|---|---|---|---|
| Auto Grooming | June 6 vs June 7 | 27.15 | 1 | < 0.001 |
| | June 6 vs June 8 | 0.847 | 1 | 0.357 |
| | June 7 vs June 8 | 5.48 | 1 | 0.019 |
| Auto Scratching | June 6 vs June 7 | 9.93 | 1 | 0.0016 |
| | June 6 vs June 8 | 19.38 | 1 | < 0.001 |
| | June 7 vs June 8 | 34.61 | 1 | < 0.001 |
| Examining Genetalia | June 6 vs June 7 | 161.76 | 1 | < 0.001 |
| | June 6 vs June 8 | N/A | N/A | 0.089 (Fishers exact) |
| | June 7 vs June 8 | 51.34 | 1 | < 0.001 |
| Hyper-vigilance | June 6 vs June 7 | 4.02 | 1 | 0.045 |
| | June 6 vs June 8 | 35.86 | 1 | < 0.001 |
| | June 7 vs June 8 | 34.05 | 1 | < 0.001 |
| Nest Construction | June 6 vs June 7 | 136.26 | 1 | < 0.001 |
| | June 6 vs June 8 | 7.70 | 1 | 0.006 |
| | June 7 vs June 8 | 66.96 | 1 | < 0.001 |

**Table S4**. Fisher's Exact test, comparing the frequency of behaviors across observation days (Day before birth: June 6; Day of birth: June 7; Day after birth: June 8) in a mother aye-aye (n=1). Pairwise comparison for those anxiety-related behaviors with expected values of < 5. Significance adjusted for Bonferroni correction (α = 0.0167)

| Behavior | Comparison | p-value |
|---|---|---|
| Nudging Infant | June 6 vs June 7 | N/A, no behaviors |
| | June 6 vs June 8 | < 0.001 |
| | June 7 vs June 8 | < 0.001 |
| Pacing | June 6 vs June 7 | 0.032 |
| | June 6 vs June 8 | 1 |
| | June 7 vs June 8 | 0.327 |
| Shaking | June 6 vs June 7 | 0.185 |
| | June 6 vs June 8 | 0.596 |
| | June 7 vs June 8 | N/A, no behaviors |
| Yawning | June 6 vs June 7 | 0.556 |
| | June 6 vs June 8 | 1 |
| | June 7 vs June 8 | N/A, no behaviors |

**Table S5.** Kruskal Wallis test analyzing the distribution of the duration of anxiety-related behavior bouts across three days peripartum (Day before birth: June 6; Day of birth: June 7; Day after birth: June 8) in a mother aye-aye (n=1). Behaviors measured include: auto-grooming, auto-scratching, and examining genitalia. Data presented with mean ± standard deviation and evaluated at ($\alpha = 0.05$).

| Behavior | Duration of Behaviors | | | p-value |
|---|---|---|---|---|
| | June 6 | June 7 | June 8 | |
| Auto-grooming | 24.09 ± 20.92 | 26.03 ± 28.08 | 33.40 ± 23.16 | 0.262 |
| Auto-scratching | 11.34 ± 11.77 | 8.43 ± 9.38 | 10.50 ± 2.51 | 0.032 |
| Examining Genitalia | 13.5 ± 11.99 | 20.13 ± 32.28 | 0, no behaviors recorded | 0.285 |
